# Single sample pathway analysis in metabolomics: performance evaluation and application

**DOI:** 10.1101/2022.04.11.487976

**Authors:** Cecilia Wieder, Rachel PJ Lai, Timothy Ebbels

## Abstract

Single sample pathway analysis (ssPA) transforms molecular level omics data to the pathway level, enabling the discovery of patient-specific pathway signatures. Compared to conventional pathway analysis, ssPA overcomes the limitations by enabling multi-group comparisons, alongside facilitating numerous downstream analyses such as pathway-based machine learning. While in transcriptomics ssPA is a widely used technique, there is little literature evaluating its suitability for metabolomics. Here we provide a thorough benchmark of established ssPA methods (ssGSEA, GSVA, SVD (PLAGE), and z-score) using semi-synthetic metabolomics data, alongside the evaluation of two novel methods we propose: ssClustPA and kPCA. While GSEA-based and z-score methods outperformed the others in terms of recall, clustering/dimensionality reduction-based methods provided higher precision at moderate-to-high effect sizes. A case study applying ssPA to inflammatory bowel disease demonstrates how these methods yield a much richer depth of interpretation than conventional approaches, for example by clustering pathway scores to visualise a pathway-based patient subtype-specific correlation network. We also developed the sspa python package (freely available at https://pypi.org/project/sspa/), providing implementations of all the methods benchmarked in this study. This work underscores the value ssPA methods can add to metabolomic studies and provides a useful reference for those wishing to apply ssPA methods to metabolomics data.

**Author summary:** Pathway analysis is a computational method used to draw insights from omics data by identifying groups of molecules (biological pathways) which are important in a study. Single-sample pathway analysis is based on the same principles as conventional pathway analysis but allows researchers to compute the enrichment of pathways at an individual-sample level. This enables pathway-based patient stratification and facilitates a multitude of downstream analyses based on pathways, such as machine learning, or pathway-based visualization which would not be possible using conventional approaches. In this work we investigated the application of single-sample pathway analysis to metabolomics data, a field in which it has not been widely used to date. We first evaluated the most popular methods for single-sample pathway analysis using simulated metabolomics data, as well as two novel methods we developed. Following these tests, we used metabolomics data from inflammatory bowel disease patients to demonstrate how single-sample pathway analysis methods can be used to infer novel pathway-based insights with the advantage of discriminating between disease subtypes. Overall, our analysis provides readers with information on the performance of single-sample pathway analysis methods, whilst highlighting the potential of such methods in metabolomics research.

## Introduction

As metabolomics continues to gain traction in various fields of research, interpretation of the results of such studies remains of paramount importance. This is particularly key in untargeted experiments, in which as many compounds as possible are assayed in order to build a comprehensive metabolic profile of the phenotype under study [1]. Such a study does not usually begin with a specific hypothesis, but seeks to develop and refine hypotheses through downstream analysis of the data. A typical metabolomics analysis workflow usually involves identifying a subset of compounds of interest in relation to the study objective, which can be achieved in various ways not limited to statistical association testing, multivariate statistical approaches, and machine learning (classification and regression) [2]. Once compounds of interest are identified, the next step often involves placing these in a biological context. Pathway analysis (PA) is perhaps the most well-known computational approach for doing so, providing users with a list of pathways considered enriched in the condition of interest (e.g. disease vs. healthy) [3]. These pathways are curated using both manual and computational approaches and deposited in databases, representing sets of biochemical reactions that collectively perform a specific function [4].

Conventional PA approaches widely used across omics data types (genomics, transcriptomics, proteomics, and metabolomics, etc) commonly focus on a two-group analysis, seeking to identify significantly impacted pathways between the study groups. These methods can be broadly classified into three main categories: over-representation analysis (ORA) [5], functional class scoring approaches such as GSEA [6], and network topology based approaches [7]. While there exists a wealth of literature describing and evaluating these approaches in transcriptomics [3,7,8], the development and application of these methods is only at the early exploration phase in the metabolomics context [9–11]. Despite the popularity and success of these methods, there are several use cases in which they are unsuitable: i) studies in which there is more than a single contrast (or continuous outcome) to be analysed, and ii) when the aim is to estimate the importance of a pathway at an individual sample level, rather than across experimental groups as a whole.

Single-sample PA (ssPA) refers to methods used to compute a score representing the enrichment level of each pathway for each individual sample in a study [12]. Another way to conceptualise ssPA is that it can be used to transform individual molecular-level data to the pathway level (Fig. 1a). For example, the metabolite abundance matrix *X_n×m_*, with *n* rows representing samples and *m* columns representing metabolites, could be transformed to the matrix *Y_n×p_* with rows representing identical samples but columns instead representing pathways (Fig. 1b). This principle of transforming a data matrix from the individual molecular measurements to the pathway space is generalisable to any type of omics data, including metabolomics. By calculating pathway scores for each sample within a dataset, the concept of ssPA overcomes the limitations of conventional PA methods, allowing for multi-group PA and enabling PA at the individual sample level.

**Fig 1:**
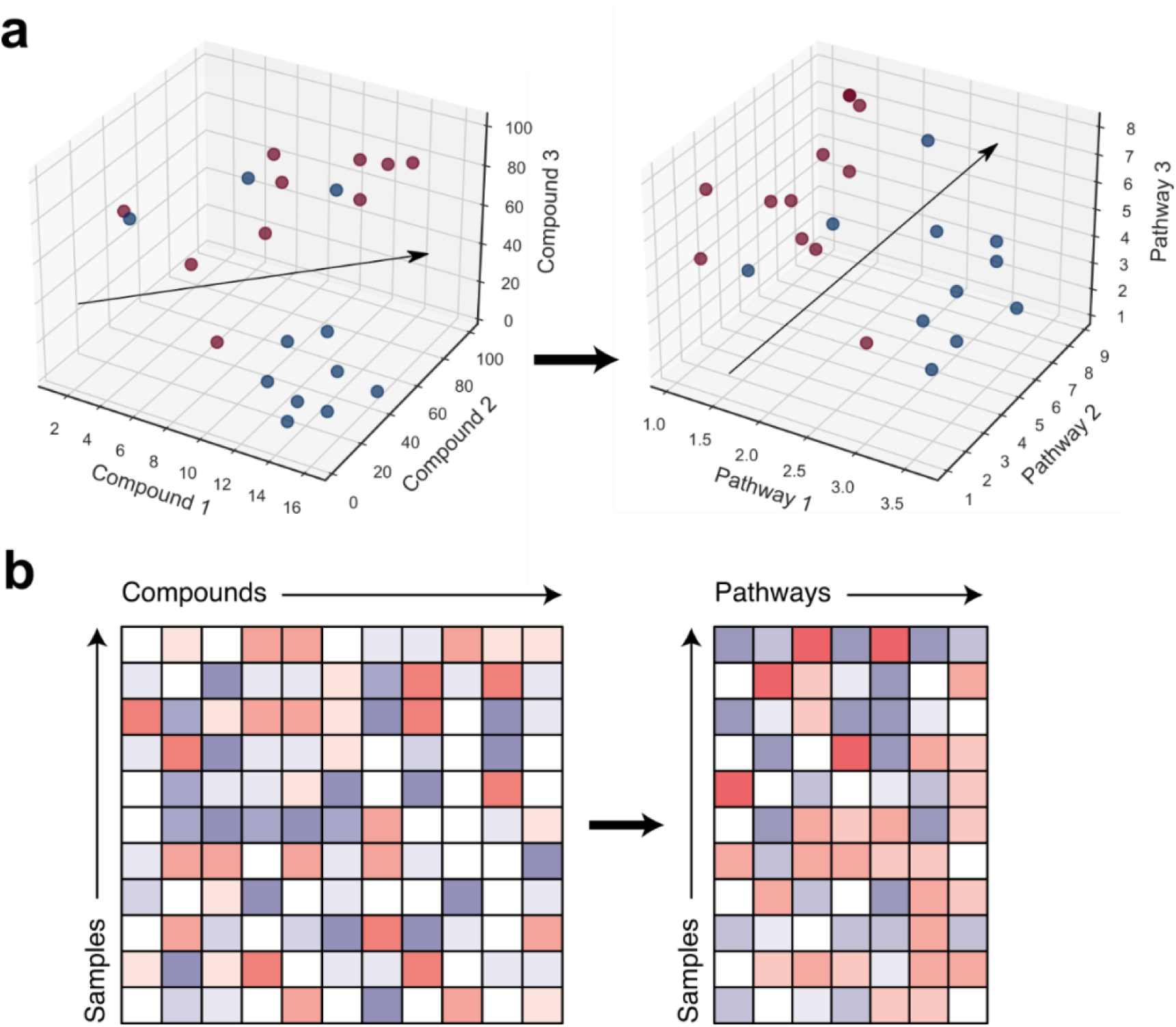
Schematic representation of single sample pathway analysis. (a) Transformation of omics data from the metabolite space (left) to the pathway space (right). Data point colours represent sample groups. (b) Left heatmap represents original omics data matrix (metabolomics) *X_n×m_*, and right heatmap represents ssPA-transformed pathway level omics data matrix *Y_n×p_*. Both matrices contain rows representing the same samples, but ssPA transforms the columns of the original matrix representing individual molecular measurements (in this example metabolites) into pathways.

Current ssPA methods can be broadly categorised into three main groups: dimensionality reduction (DR)-based [13], GSEA-based [14,15], and z-score-based [16]. The earliest example of ssPA can be attributed to Tomfohr et al. [13], who developed the PLAGE method for calculating pathway scores using singular value decomposition (SVD). ssGSEA and GSVA are two similar ssPA methods based on the Kolmogorov-Smirnov like random walk statistic proposed by Subramanian et al., 2005 [6] for conventional GSEA. The z-score method developed by Lee et al., 2008 [16] calculates pathway scores based on a normalised z-score. Full details of these methods can be found in the original articles [13–16].

By generating sample-wise pathway scores, ssPA enables numerous downstream analyses to be performed that cannot be achieved using conventional PA approaches. At the most basic level, multi-group comparisons can be made using the pathway scores, for example using statistical association testing to determine pathways which differ significantly between groups [15]. This can be useful in studies where there are more than two treatment groups or disease subtypes. Another prominent example of the utility of ssPA methods is that they enable application of machine learning or multivariate statistical methods to pathway level data [17]. Patients can be classified based on their pathway scores, rather than metabolite-level measurements, which in some cases has been demonstrated to improve classification performance [16]. Predictive models based built in the pathway space are also known to be more robust to noise than those constructed from individual molecular measurements [18]. Pathway scores further enable a number of pathway-based visualisation options, such as plotting the scores of two pathways against each other to discriminate between disease subtypes [19], or using them to generate hierarchical clustering heatmaps [13]. An emerging use-case of ssPA is in multi-omics data integration, in which pathway scores calculated for each omics layer can be combined and analysed concurrently [20].

Although most ssPA methods are applicable to most omics datatypes, this will not necessarily equate to their consistent performance across omics. Indeed, the most well-known ssPA methods have all been developed for transcriptomic data analysis. Other omics datatypes such as metabolomics differ greatly in composition and statistical properties from transcriptomic data. The most obvious difference lies in the number of metabolites versus the number of genes/transcripts profiled in metabolomics and transcriptomics respectively, which has a direct effect on pathway coverage (the proportion of entities in a given pathway that have been assayed). Another key difference is the uncertainty in compound identification in metabolomics which remains one of the field’s greatest challenges [21], an issue that is far less prominent in transcriptomics and proteomics. Despite the potential of ssPA methods for metabolomics, to date there have been no studies investigating the efficacy of these methods when applied to metabolite datasets. The purpose of the present research, therefore, is to perform a comprehensive benchmark of ssPA methods applied to both real (experimental) and semi-synthetic metabolomics datasets. We begin by evaluating the performance of four well-known ssPA methods, namely SVD (PLAGE), ssGSEA, GSVA, and z-score using semi-synthetic data, and then demonstrate using experimental data the value ssPA can add to a metabolomics study. We also propose two novel methods for ssPA, ssClustPA and kPCA which are available freely through the `sspa` package. This is the first study to benchmark ssPA methods on metabolomics data, offering important insights into their suitability and potential applications in metabolomics research.

## Methods

### Datasets used

In the benchmarking section of this work, the Su et al., 2020 [22] COVID19 metabolomics mass spectrometry (MS) dataset has been used, serving as a basis for semi-synthetic simulated data creation. This data is part of a multi-omics study of COVID severity, and contains patients from two groups: COVID (of varying levels of severity) (n=130) and non-COVID (n=133), from which blood plasma samples were obtained during the first week of infection after diagnosis.

For the application section, we used inflammatory bowel disease (IBD) MS metabolomics data from Lloyd-Price et al., 2019 [23]. This data derives from a longitudinal study of multi-omics in IBD, where metabolomics analysis was performed on stool samples collected over a one-year period. We used samples obtained across all timepoints (weeks 0-52). The study contains two experimental groups: IBD (Crohn’s disease (CD, n=265) and ulcerative colitis (UC, n=146)) and non-IBD (n=135).

Both datasets are derived from untargeted mass spectrometry, see Table 1. Datasets were post-processed in the same manner: missing value imputation using SVD, probabilistic quotient normalisation, *log*_2_ transformation, and standardisation of each variable (*μ* = 0 and *σ* = 1).

**Table 1:**
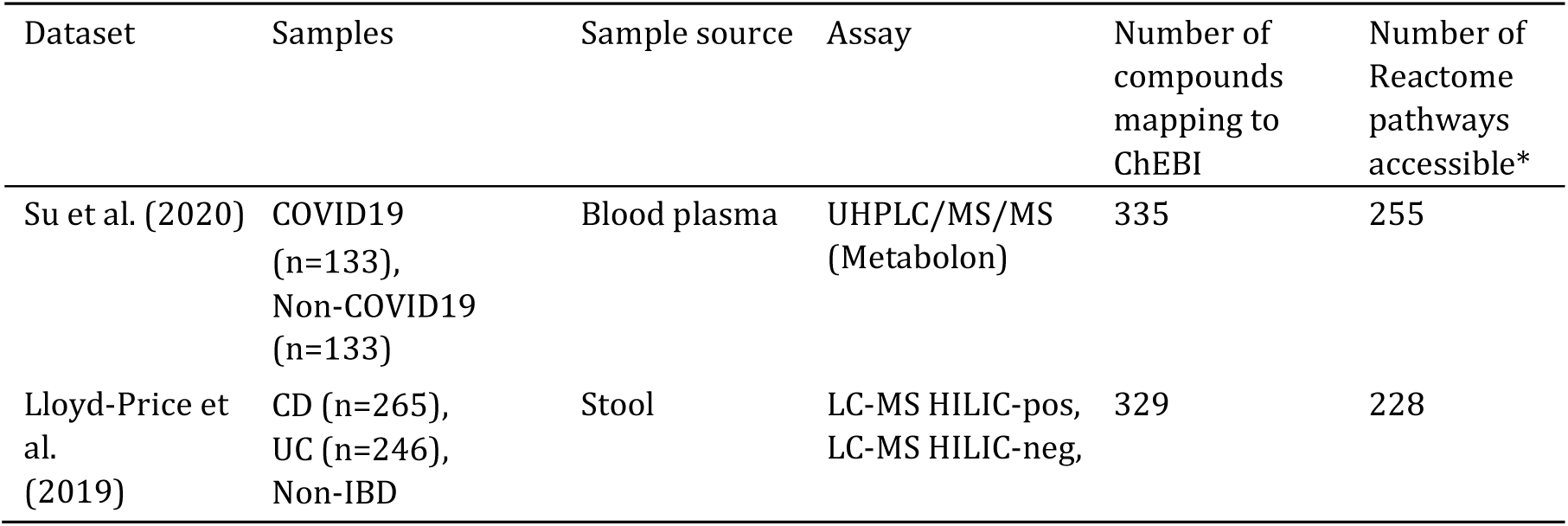
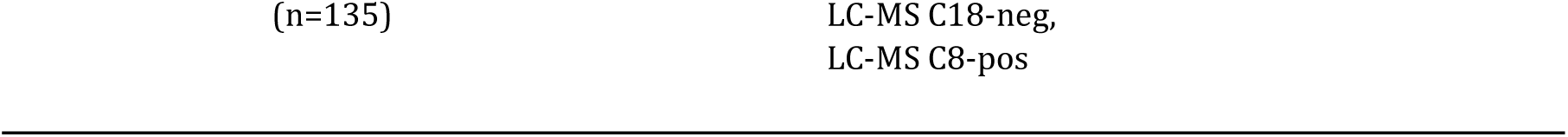
Summary of datasets used. *Number of Reactome pathways accessible corresponds to the number of pathways in each dataset which contained at least two compounds assayed in the metabolomics data.

### Pathway definitions and non-redundant pathway set

Pathways from Reactome release 76 were downloaded from https://reactome.org/download-data. The `ChEBI2Reactome_All_Levels.txt` file was used and filtered for human pathways only.

As part of the benchmarking procedure a non-redundant set of Reactome pathways *N* = {*p*_1_, *p*_2_, …, *p*_*N*_} was created along with the associated set of all metabolites in these pathways *M* = {*m*_1_, *m*_2_, … *m*_*M*_} = *p*_1_ ∪ *p*_2_ ∪…∪ *p*_*N*_. *N* and *M* were obtained by iterating through the list of pathways *P* (*k*=255 with at least 2 compounds present profiled in the [NO_PRINTED_FORM]COVID dataset, in original order) and adding pathway *p*_*i*_ to *N* if there was no overlap with the current non-redundant set, |*p*_*i*_ ∩ *M*| = 0. The resulting set contained a total of 18 pathways with no overlapping compounds, with coverage ranging from 2 to 6 compounds per pathway. We note that other non-redundant pathway sets are possible but do not expect results to be strongly dependent on the exact set used.

### Creation of semi-synthetic metabolomics data

The simulations in this work are based on semi-synthetic datasets created Su et al., 2020using the COVID dataset. The use of a “permutation and spike” procedure allows the original signal in the dataset to be erased and replaced with known pathway signals, while preserving the underlying distributions (both joint and marginal) and heterogeneity of the experimental data. This is advantageous compared to generating fully synthetic data such as by random sampling from a Gaussian distribution, as it preserves the complex biological relationships occurring in omics data, which will influence method performance. A limitation of our approach is that the underlying distribution of a particular dataset may differ from that of other datasets. Despite this limitation, we believe this approach reflects more accurately the level of complexity found in real data compared to fully-synthetic simulation approaches.

Let *X*_*i*,*j*_ denote the original *log*_2_-transformed data matrix composed of *n* samples and *m* metabolites. *X* contains two groups of samples, *i* ∈ *A*, representing the control group and *M* ∈ *N*, representing the disease group. For simplicity we have simulated a two-group design, but this simulation setup can be extended to multiple groups, continuous outcomes or more complex experimental designs. To remove the original signal in the data, the class labels of each sample *M* or *N* were randomly shuffled at each realisation of the simulated data. Following this, *k* pathways {*p*_1_, *p*_2_, … , *p*_*k*_} were chosen at random to be “enriched”, corresponding to metabolites *M*_*k*_ = {*p*_1_ ∪ *p*_2_ ∪…∪ *p*_*k*_}. The abundance values samples in group *B* were altered (by adding a constant *α* to the original log space values) for compounds in set *M*. Note this represents a multiplicative (fold change) effect in the original non-log space. The values for group *A* were left unchanged. The value of *α* represents the strength of the enrichment of the compounds within *p*_*i*_. In our experiments we enriched *k* = 3 pathways. The simulated data matrix *Y* can thus be expressed as:

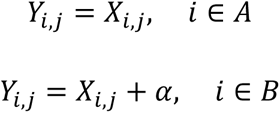

where *j* ∈ *M*_*k*_ and *α* ∈ [0,1].

### ssPA implementation details

Details of the Python/R packages used for each of the methods benchmarked in this work are given in Table S1. The `ssPA` package is available to download from the Python Package Index at https://pypi.org/project/sspa/, and the source code is freely available at https://github.com/cwieder/py-ssPA.

### ssClustPA method

Fig. S1 shows an overview of ssClustPA and kPCA methods. The ssClustPA method is outlined as follows:

1. For each of the *P* pathways *p*_*k*_ = {*m*_1_, *m*_2_, … , *m*_*M k*_ }, create a pathway matrix *Z*_*k*_ composed of the abundance data corresponding to the pathway, *Z*_*ijk*_ = *X*_*ij*_ where *m*_*j*_ ∈ *p*_*k*_. Here there are *M_k_* metabolites in pathway *k*.
2. On each pathway matrix *Z*_*k*_, perform k-means clustering based on Euclidean distance with `2` clusters.
3. Obtain the coordinates of the cluster centroids *c*_*g*_ in the *m*-dimensional space defined by *Z*_*k*_.
4. As Euclidean distances cannot capture the position of a sample data point relative to the cluster centroids, ssClustPA uses the vector between the two cluster centroids and projects the sample datapoints onto this vector to produce pathway scores with directionality relative to the cluster centroids. For each pathway obtain the unit column vector between the two cluster centroids *u* = *c*_1_ − *c*_2_. The dimension of *u* will correspond to the number of metabolites in the dataset mapping to that pathway, *M_k_*.
5. Project the pathway matrix *Z*_*k*_ (dimension n × M_k_) onto *u* (dimension M_k_ × 1). The projected values can be used as pathway scores *A*_*k*_ (Eqn. 1).

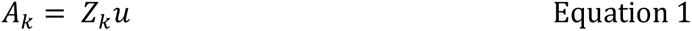
6. Repeat steps 4-5 for each pathway matrix *Z*_*k*_ and concatenate the resulting vectors *A*_*k*_ horizontally to produce the pathway score matrix *A*_*n*×*P*_.

### kPCA method

The kPCA method is outlined as follows:

1. For each of the *P* pathways *p*_*k*_ = {*m*_1_, *m*_2_, … , *m_M k_*}, create a pathway matrix *Z*_*k*_ composed of the abundance data corresponding to the pathway, *Z*_*ijk*_ = *X*_*ij*_ where *m*_*j*_ ∈ *p*_*k*_.
2. For each pathway matrix *Z*_*k*_, perform kernel PCA [24] with a radial basis function (RBF) kernel. The kernel width parameter *γ* is set to a default of 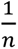.
3. The scores of the first principal component (PC1) can directly be used as scores *A*_*k*_.
4. Repeat step 2 for each pathway matrix *Z*_*k*_ and concatenate the resulting vectors *A*_*k*_ horizontally to produce the pathway score matrix *A*_*n*×*P*_.

### Benchmarking details

#### Pathway overlap

Overlap between pathway compounds was calculated using the Szymkiewicz-Simpson Overlap Coefficent (OC). An OC of 0 indicates there is no overlap between set A and set B, whereas an overlap of 1 indicates that the smaller set is a subset of the larger set.

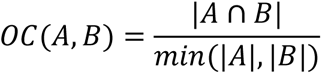

We used the OC as an alternative to the Jaccard Index (JI) as it is more sensitive to overlapping compounds, i.e. the OC will return a value of 1 when the compounds in a pathway are all present in a larger pathway, whereas the JI will only equal 1 if two sets are identical.

#### Performance metrics

In each of the simulations in this work, *k=3* randomly selected pathways *E* = {*p*_1_, *p*_2_, *p*_3_} were defined as enriched. In the effect size simulations, all metabolites in the set *E* had a constant *α* ∈ [0,1] added to the value of those metabolites in the disease class. Adding a constant of 1 to a metabolite in the log space changes its abundance value by a fold change (FC) of 2 (*log*_2_*FC* = 1). In the signal strength simulation, varying percentages *s* ∈ [0,100] of randomly selected metabolites in *E* had a constant *α* = 1 added to them, again only in the disease class.

Pathway scoring using ssPA was performed on the simulated data abundance matrix, and a series of independent two-sample t-tests between simulated disease and control groups were used to determine the significance of each pathway. The Benjamini Hochberg false discovery rate correction was applied and pathways with q ≤ 0.05 were considered significantly enriched. Of these pathways, those that were members of *E* or those with an OC ≥ θ were considered true positives. We tested two OC thresholds of θ ∈ {0.25,0.5}. We use the OC to define true positives as pathways with overlapping compounds are likely to also be enriched. Pathways with q ≥ 0.05 were considered true negatives if the pathway is not a member of *M* and has OC ≤ θ. Those pathways with q ≥ 0.05 that are members of *E* or have OC ≥ θ are considered false negatives.

For a given pathway *p*:

*p* is a true positive (TP) if *p* ∈ *E* or OC(E, p) ≥ θ, and *q* ≤ 0.05
*p* is a true negative (TN) if *p* ∉ *E* or OC(E, p) ≤ θ, and *q* ≥ 0.05
*p* is a false positive (FP) if *p* ∉ *E* or OC(E, p) ≤ θ, and *q* ≤ 0.05
*p* is a false negative (FN) if *p* ∈ *E* or OC(E, p) ≥ θ, and *q* ≥ 0.05

The `sklearn.metrics` functions were used to calculate recall, precision, and area under the receiver-operating curve (AUC). All simulations were repeated 200 times, with randomly selected enriched pathways and semi-synthetic data randomly permuted at each realisation.

#### Method runtime profiling

Runtime profiling was repeated 10 times for each method and average results are reported. The Python libraries `cProfiler` and `line-profiler` were used to determine the wall time of each method.

### IBD application

The kPCA method was used to demonstrate ssPA applied to the IBD metabolomics dataset.

#### Hierarchical clustering

kPCA pathway scores were standardised prior to clustering (μ = 0 and σ = 1). Hierarchical clustering was performed using the `scipy cluster.hierarchy` function with Euclidean distance and Ward linkage parameters. The `maxclust` parameter was set to 2 in order to cut the tree at a depth of 2 branches. The adjusted Rand index was calculated using the `metrics.adjusted_rand_score` function, with the original and predicted cluster sample labels as input.

#### Pathway score correlation network

kPCA pathway scores derived from the IBD dataset were ranked by t-test p-values (testing for differences between IBD and non-IBD groups) and the top 50 pathways were used to produce a hierarchical clustering as detailed above. Pathway IDs from the resulting clusters were extracted and used to build the network. A Spearman rank correlation matrix was produced from the pathway scores. `networkx` was used to create a pathway-pathway network, with nodes representing pathways and edges representing correlation between the pathway scores. Cytoscape was used to visualise the network using an edge-weighted spring embedded layout.

The COVID pathway correlation network was created in the same way as the IBD network, using the top 30 pathways (ranked by t-test p-values) for visual clarity. Each node is coloured by the average pathway score of samples in each COVID WHO severity level (0, 1-2, 3-4, 5-7).

## Results

### Performance evaluation of ssPA methods

We first carry out a comprehensive benchmarking of ssPA methods applied to semi-synthetic metabolomics datasets generated based on the Su et al. COVID dataset [22], see Methods. All pathways used throughout this work were obtained from the Reactome pathway database [25]. Where applicable, we also compare ssPA results to those obtainable using conventional PA methods ORA and GSEA.

#### Outline of simulation procedure

In order to calculate various performance metrics (i.e. precision and recall), it is essential to know the identity of truly perturbed “enriched” pathways. Here we use the terms “positively enriched” or “negatively enriched” to refer to pathways that are significantly perturbed with respect to the control group, depending on the directionality of the effect. In experimental datasets it remains challenging to distinguish between true and false positive enriched pathways, so instead we use semi-synthetic data to accomplish this task. As detailed in the methods, we remove the original signal from an experimental untargeted metabolomics dataset of COVID patients [22]and replace it with artificial known pathway signals (enriched pathways) in one of two study groups. Using this simulation procedure, we can vary the strength of the enrichment of a pathway and the number of differentially abundant metabolites in the pathway.

#### ssClustPA and kPCA: novel pathway scoring approaches

We propose two novel methods based on unsupervised machine learning concepts for generating pathway scores (see Methods for further details). ssClustPA makes use of k-means clustering and for each pathway projects the original datapoints onto the unit vector between two cluster centroids to yield pathway scores. The kPCA method uses a radial basis function kernel to model the distribution of data for the pathway, and is particularly advantageous when the underlying manifold of the data is non-linear. The first principal component scores are used as pathway scores. Both methods are applicable to any omics datatype.

#### The importance of pathway overlap in method evaluation

Due to the nature of biological pathways and how metabolic networks are connected as a whole, defining precise pathway boundaries remains a challenge in pathway curation. It is therefore expected that all pathway databases will contain some degree of redundancy, meaning that not all molecules will be unique to a single pathway. If a pathway *p* is significantly enriched, it is likely that other pathways that contain overlapping molecules with *p* will also be considered enriched to varying degrees. When performing benchmarking on redundant pathway sets, one must therefore decide the level of pathway overlap that constitutes a true positive pathway.

A simple example of the effect of pathway overlap is demonstrated in Fig 2. Here we performed a simulation in which a single randomly selected pathway “R-HSA-1483206” (Glycerophospholipid biosynthesis) was enriched at effect size *α* = 1. Using the pathway scores derived from each ssPA method, a series of independent two-sided t-tests were performed for each of the 255 Reactome pathways, testing for significant differences in mean pathway scores between the two study groups. The resulting pathway p-values are compared to the overlap coefficient (OC) of each pathway with pathway R-HSA-1483206 in Fig. 2. We can observe that regardless of the ssPA method, there is a clear correlation between pathway overlap and p-value, with pathways with lower overlap tending to have lower p-values. Importantly, if an arbitrary p-value threshold e.g. p ≤ 0.05 was used to select significant pathways in this manner, R-HSA-1483206 and a number of additional pathways would be considered enriched. It is important to note this effect is not only observed with ssPA methods, but also in conventional methods such as GSEA (Fig. 2).

**Fig 2:**
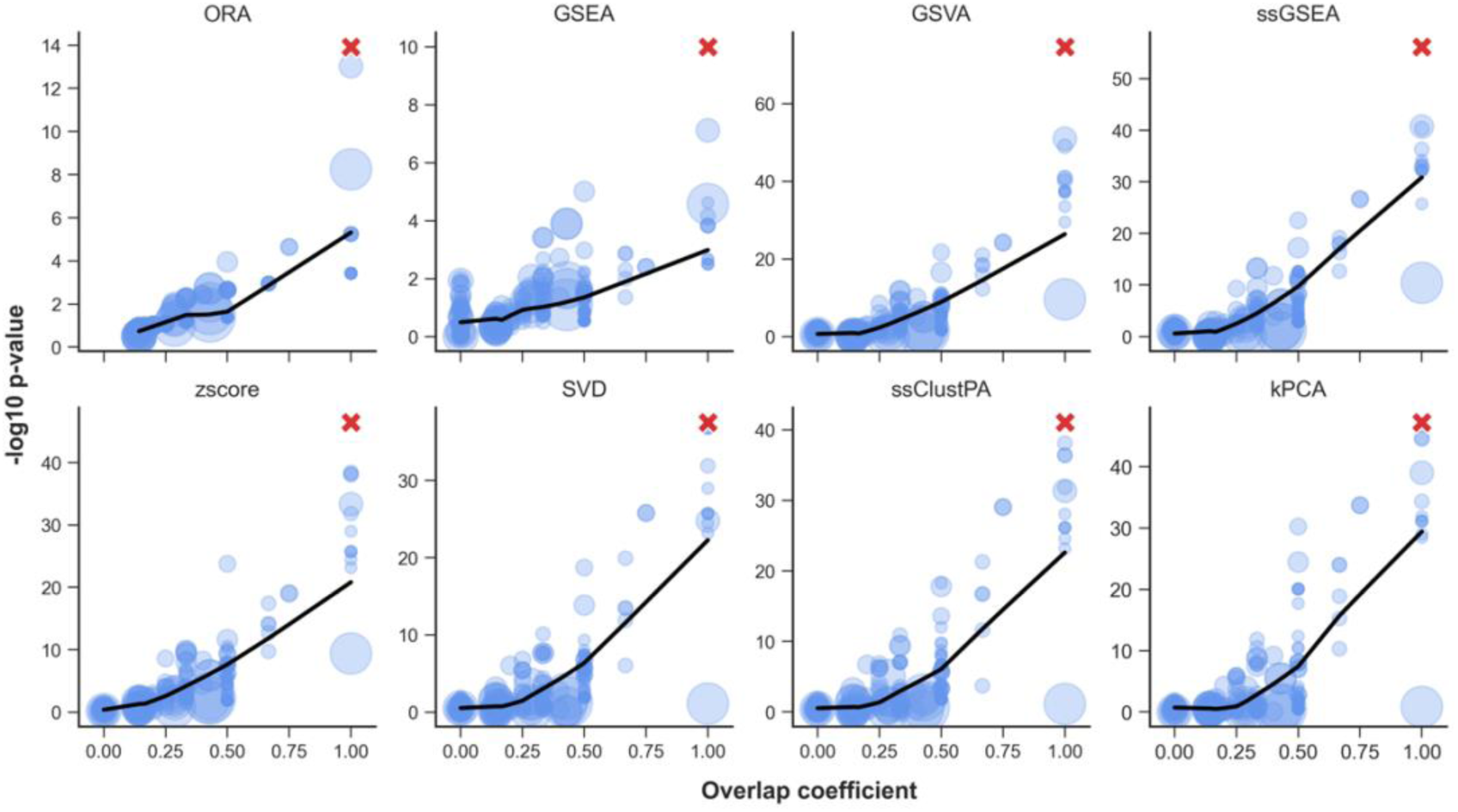
The relationship between pathway overlap and significance. The artificially enriched pathway (R-HSA-1483206) is represented by a red cross. Overlap coefficient values of each pathway with R-HSA-1483206 are shown on the x-axis. t-test p-values of pathway scores (testing for significant difference in mean pathway score between disease and control group) for each pathway are shown on the y-axis (-log10 scale). Note, for GSEA the p-values are calculated using the original GSEA permutation procedure, and for ORA, p-values are calculated using the Fisher’s exact test. A LOWESS regression line is shown in black. Point size corresponds to the coverage of each pathway (i.e. the number of compounds in the pathway which were present in the dataset).

Due to this overlap effect, and to simplify the procedure, we first perform benchmarking on a small subset of non-redundant pathways from the Reactome pathway database (*k*=18). We then repeat the performance evaluation on the full set of redundant Reactome pathways (*k*=255 with sufficient coverage in the dataset), which although containing overlapping pathways, is the most realistic scenario.

#### Performance evaluation of ssPA methods

In this section we evaluated the performance of the SVD (PLAGE), ssGSEA, GSVA, z-score, ssClustPA, and kPCA methods using the semi-synthetic data. These are categorised as GSEA-based (ssGSEA and GSVA) and dimensionality reduction (DR)/clustering-based (SVD, ssClustPA, and kPCA). We also included comparison to non-ssPA methods ORA and GSEA where possible.

Beginning with the non-redundant pathway set, in which each of the pathways contains unique compounds, three random pathways were artificially enriched and performance metrics were computed and averaged over 200 iterations (Fig 3) across a range of increasing effect sizes. The GSEA-based methods GSVA and ssGSEA appear to outperform the other methods in terms of recall and AUC at low to moderate effect sizes (0.2-0.6). At higher effect sizes of 0.8-1 (corresponding to *FC* ≈ 2), all methods appear to perform equally well with average recall, precision and AUC values > 0.9.

**Fig 3:**
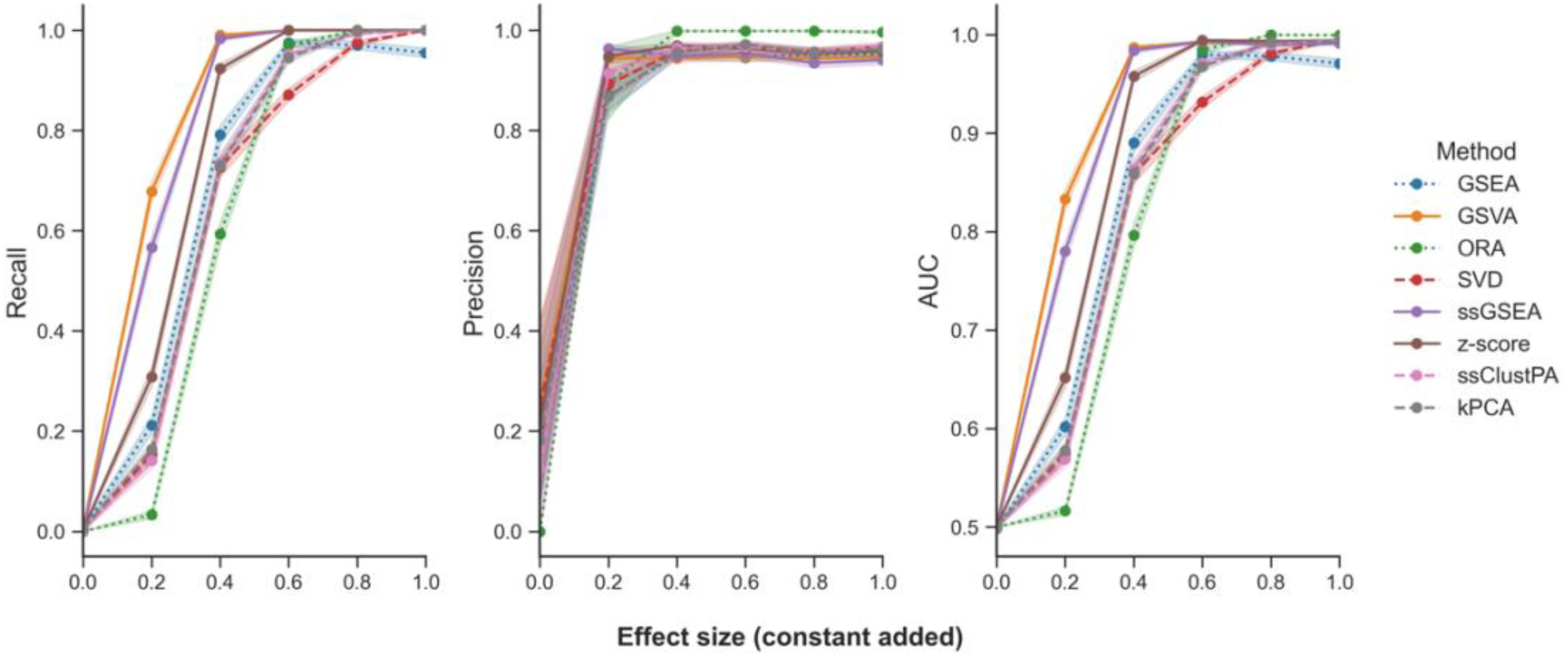
Performance of ssPA methods using a non-redundant pathway set based on 3 randomly selected enriched pathways. For each panel Recall, Precision, and AUC, the x-axis represents the effect size of the simulated pathway enrichment. Points represent the mean performance metric over 200 iterations. Shaded intervals indicate the standard error of the mean. Dotted lines represent conventional pathway analysis methods, and dashed lines represent clustering/dimensionality-reduction based methods.

We then used the full set of 255 Reactome pathways (containing redundancy) with same simulation setup to evaluate performance (Fig 4). Again, 3 random pathways are artificially enriched. As mentioned previously, when calculating performance metrics using overlapping pathways, one must define a threshold for true positive pathways. In this case, we pooled all metabolites from the simulated enriched pathways together and calculated an OC between this set and each pathway tested. Fig 4a shows the performance metrics where a positive corresponds to OC ≥ 0.25, whereas Fig 4b shows the performance metrics where a positive corresponds to OC ≥ 0.5. In both Figs. 4a and 4b, the GSEA-based methods (ssGSEA and GSVA) and z-score have the highest recall across all effect sizes. In terms of precision, regardless of the overlap threshold, conventional pathway analysis methods ORA and GSEA outperform ssPA methods at moderate-high effect sizes (0.4-1.0). Following this, DR/clustering-based methods (kPCA, ssClustPA, and SVD respectively) provide the highest precision values of the ssPA methods, with GSEA-based methods ssGSEA and GSVA as well as the z-score yielding the lowest precision values. Similar observations can be made in terms of AUC at an overlap threshold of ≥ 0.5 (Fig. 4b). The GSEA-based methods and z-score yield the highest AUC at lower effect sizes (0.2-0.4), but at higher effect sizes (0.4-1), ORA and GSEA outperform all ssPA methods, followed by the clustering/DR-based methods, and finally GSEA/z-score-based methods which have the lowest AUC values. At an overlap threshold ≥ 0.25 (Fig. 4a), these trends are more subtle in terms of AUC, but as effect sizes become larger there is a shift in performance from GSEA-based/z-score methods to clustering/DR-based methods.

**Fig 4:**
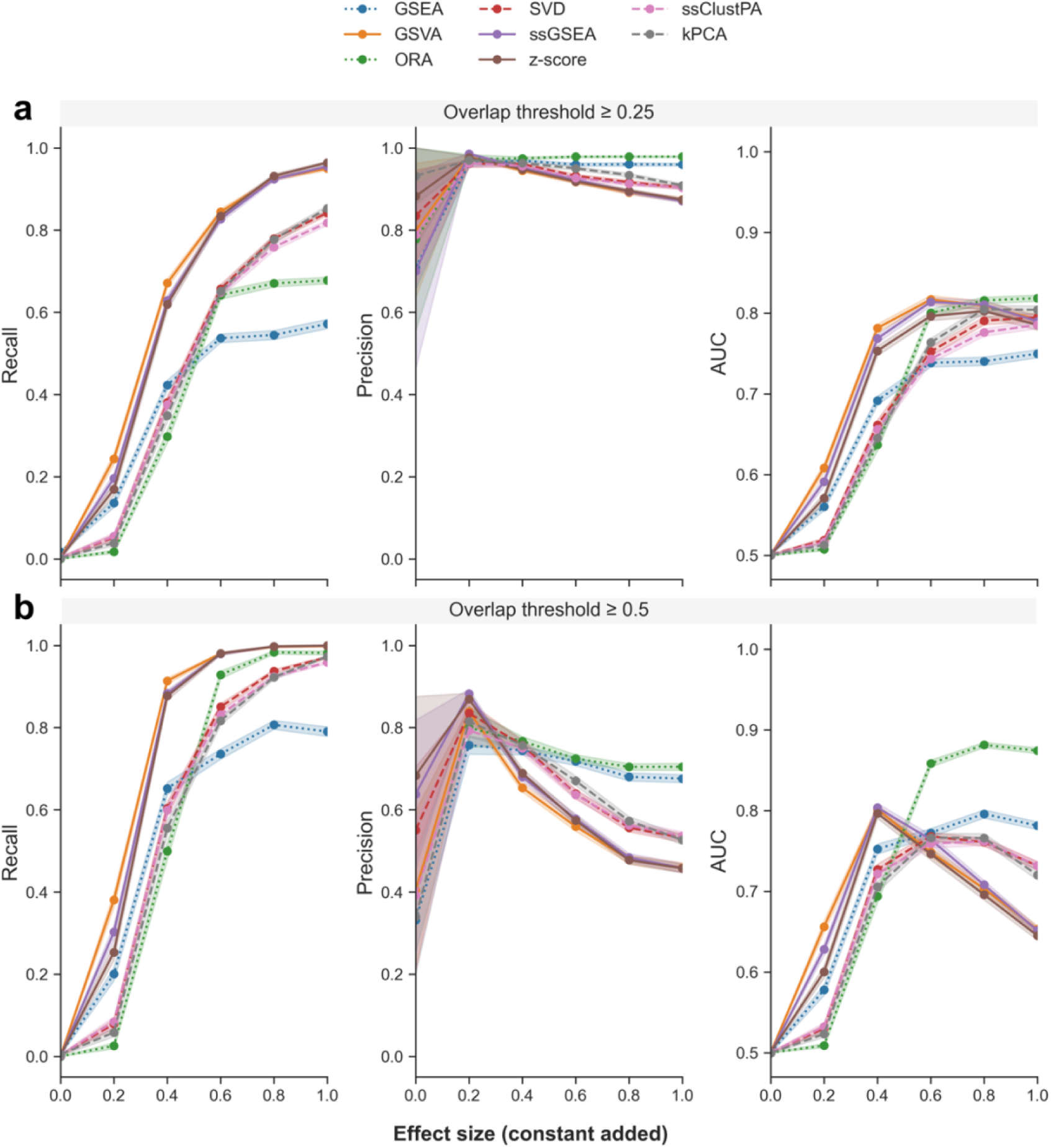
Performance of ssPA methods on the full pathway set (including redundancy) based on 3 randomly enriched pathways. (a) top panel OC ≥ 0.25, (b) bottom panel OC ≥ 0.5. All compounds in enriched pathways have identical effect size. Points show performance metrics averaged across 200 iterations. Shaded intervals represent average SEM. Dotted lines represent conventional pathway analysis methods, and dashed lines represent clustering/dimensionality-reduction based methods.

GSEA-based methods, as well as the z-score, are the poorest performers in terms of precision. The drop in precision from effect size 0.2 onwards is likely due to an increase in the number of false positives resulting from the stronger effect size which do not reach the overlap threshold to be considered true positives. There is a larger probability that overlapping pathways are detected as significantly enriched. This drop in precision is less evident when the OC for identifying true positive pathways is lower, such as in Fig. 4a, as a higher proportion of the overlapping pathways are considered true positives, resulting in fewer false positives. Compared to conventional PA methods, ssPA methods generally have improved recall, particularly at larger effect sizes. However, conventional methods tend to have a higher precision than ssPA methods, regardless of effect size. At higher effect sizes, ORA generally appears to have higher power to detect significant pathways than GSEA. Finally, we reproduced very similar results using a different semi-synthetic dataset based on data from Lloyd-Price et al., 2019 [23] (Fig. S2). The sole difference was that kPCA had higher precision than all other methods at the higher OC threshold, regardless of effect size.

#### The effect of pathway signal strength on ssPA performance

In the previous simulations, we varied the effect size of the pathway enrichment, with the abundance of all metabolites in the pathway modified to the same extent. In a real-world scenario, it is highly unlikely that all compounds in a differentially active pathway would have the same effect size. We therefore performed another simulation in which the abundance of varying proportions of randomly selected compounds in a pathway was modified while keeping the effect size at a constant of 1.0. In this scenario, we consider pathways with q ≤ 0.05 and OC ≥ 0.5 true positives and randomly enriched 3 pathways in each iteration.

As expected the performance of all methods improves as the signal strength increases (Fig. 5), aside from a small drop in precision which is caused by the increase in signal strength rendering overlapping pathways more significant, as highlighted in the previous simulation (Fig. 4). In terms of recall, all ssPA methods perform very similarly across signal strength sizes, and clearly outperform the conventional PA methods ORA and GSEA. In contrast, the conventional methods slightly outperform the rest of the ssPA methods in terms of precision, of which the dimensionality-reduction based methods (SVD, ssClustPA, and kPCA) have higher precision than GSEA/z-score-based methods. Overall, when taking into account varying levels of signal strength, and that not all molecules in an enriched pathway are likely to be significantly differentially abundant, clustering/DR-based methods appear to offer the best overall performance in terms of ssPA, particularly at moderate to high effect sizes (FC ≈≥ 1.5).

**Fig 5:**
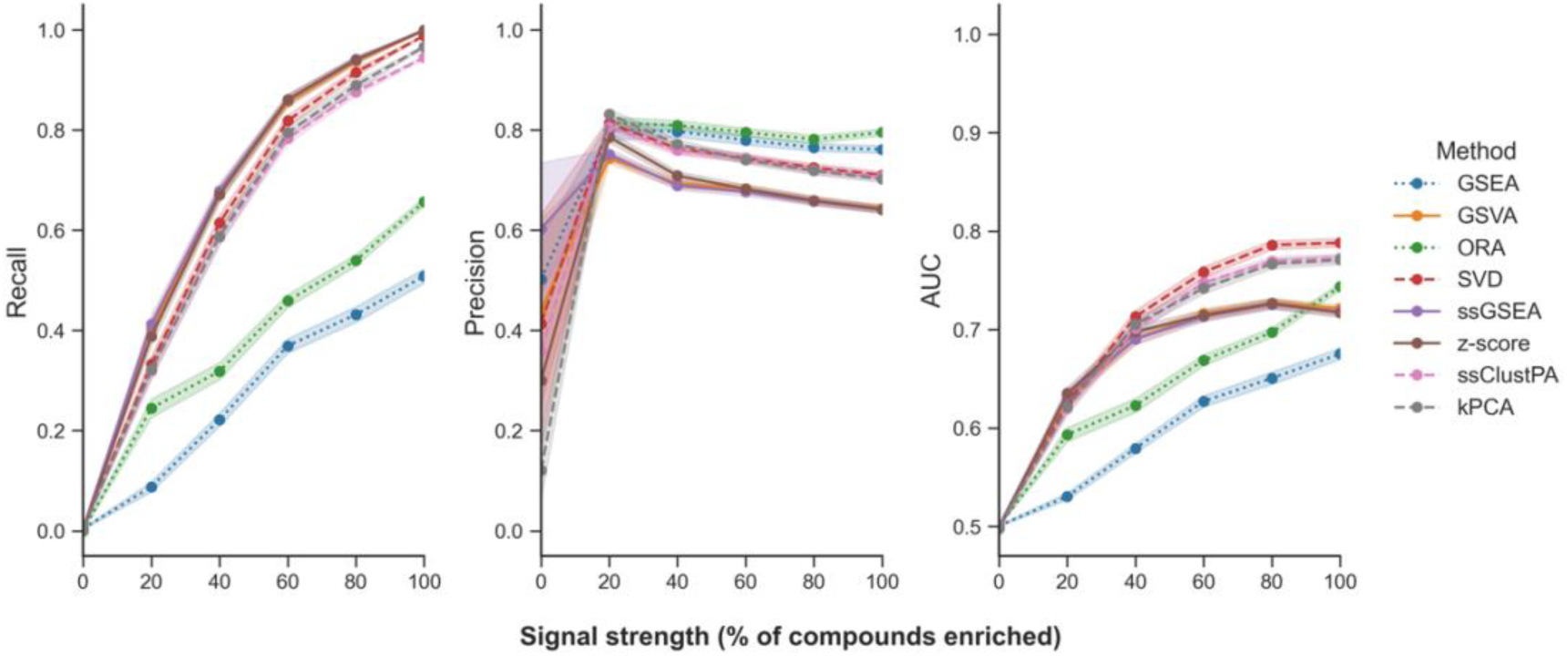
Effect of varying signal strength on ssPA method performance. Points show mean performance over 200 iterations with 3 randomly enriched pathways on the full Reactome pathway set (true positive pathways are those with an OC ≥ 0.5). Shaded intervals represent average SEM. Dotted lines represent conventional pathway analysis methods, and dashed lines represent clustering/dimensionality-reduction based methods.

#### ssPA method ability to rank enriched pathways highly

The ability of the ssPA methods to rank enriched pathways highly was investigated, again using the semi-synthetic COVID dataSu et al., 2020. Briefly, t-tests were performed on the ssPA scores for each pathway (testing for a difference between mean scores in disease and control groups) and the resulting p-values were used to rank them in ascending order. The full set of 255 Reactome pathways was used. In order to account for the fact that ORA generally tests fewer pathways than the other methods (as it only calculates p-values for pathways with at least one differentially abundant metabolite), the ranks were normalised by the total number of pathways tested using each method.

In concordance with the performance metrics calculated in the previous section, at lower effect sizes, the GSEA/z-score-based methods rank the truly enriched pathways the highest (Fig. S3). At higher effect sizes (0.8-1.0), there is very little difference in the rankings of the enriched pathways by all ssPA methods, with all methods being able to rank the 3 enriched pathways within the top 10% of pathways.

#### Method runtimes

Each ssPA method was run 10 times on a laptop with standard hardware and 16GB of RAM using the Su et al., 2020COVID dataset, which contained 263 samples (rows) and 335 compounds (columns), as well as Reactome pathways with sufficient coverage in the dataset (*k*=255). In terms of runtime, the fastest method was the z-score, followed by SVD (Table S2).

### Application of ssPA in metabolomics: a case-study of inflammatory bowel disease

The remainder of this paper focuses on the application and interpretation of ssPA in typical metabolomics data, using real experimental datasets. In order to simplify our work, we demonstrated the use of a single method, kPCA, to showcase one of our newly proposed methods, although the following application is generalisable to any ssPA method.

In order to demonstrate some of the potential use cases of ssPA methods applied to metabolomics data, we selected a different dataset to that used in the benchmarking portion of this work and made use of ssPA to help detect and interpret complex biological patterns within the data. The untargeted metabolomics data used in this case study was derived from a multi-omics study of inflammatory bowel disease (IBD) by Lloyd-Price et al., 2019. It is important to note that the group termed “non-IBD” are not necessarily healthy individuals and may still exhibit symptoms of IBD. Further details of this dataset can be found in the Methods and the original article [23].

To obtain an exploratory perspective on the strength of the biological signals in the dataset, PCA was used to visualise the data at both the metabolite and pathway level. We transformed the metabolite-level data to pathway scores using the kPCA method described earlier and compared the PCA results obtained. No strong separation between the sample classes was observed using either approach, with most samples amalgamated together in a central cluster (Fig. S4). This is consistent with findings from the original work, in which intra-individual variation in the IBD metabolome was greater than inter-individual variation [23].

In order to more rigorously quantify the clustering performance achieved using metabolites (only those present in pathways) vs. pathway scores, we ranked each of the metabolites/pathways using BH-FDR corrected t-test q-values (testing for differences between the IBD and non-IBD groups) and performed clustering after progressively adding each entity significant at q ≤ 0.05 (Fig. 6). Hierarchical clustering was used to partition the samples into groups based on the two main branches of the tree, which were compared to the original sample labels (IBD or non-IBD) using the Adjusted Rand Index (ARI). Note that negative ARI values can be obtained if the RI is less than expected by chance. This heuristic method suggested that for this particular dataset, using pathway scores for clustering generally achieves higher ARI across a range of different cut-off thresholds, as well as improved robustness to the threshold used for clustering, than by using metabolites alone. Consequently, we made use of the clustering applied to the top 50 sets of pathway scores to identify clusters of pathways that could discriminate between the IBD sample classes. Two distinct pathway clusters were evident (Fig. S5), which we visualised using Cytoscape in Fig 7.

**Fig 6:**
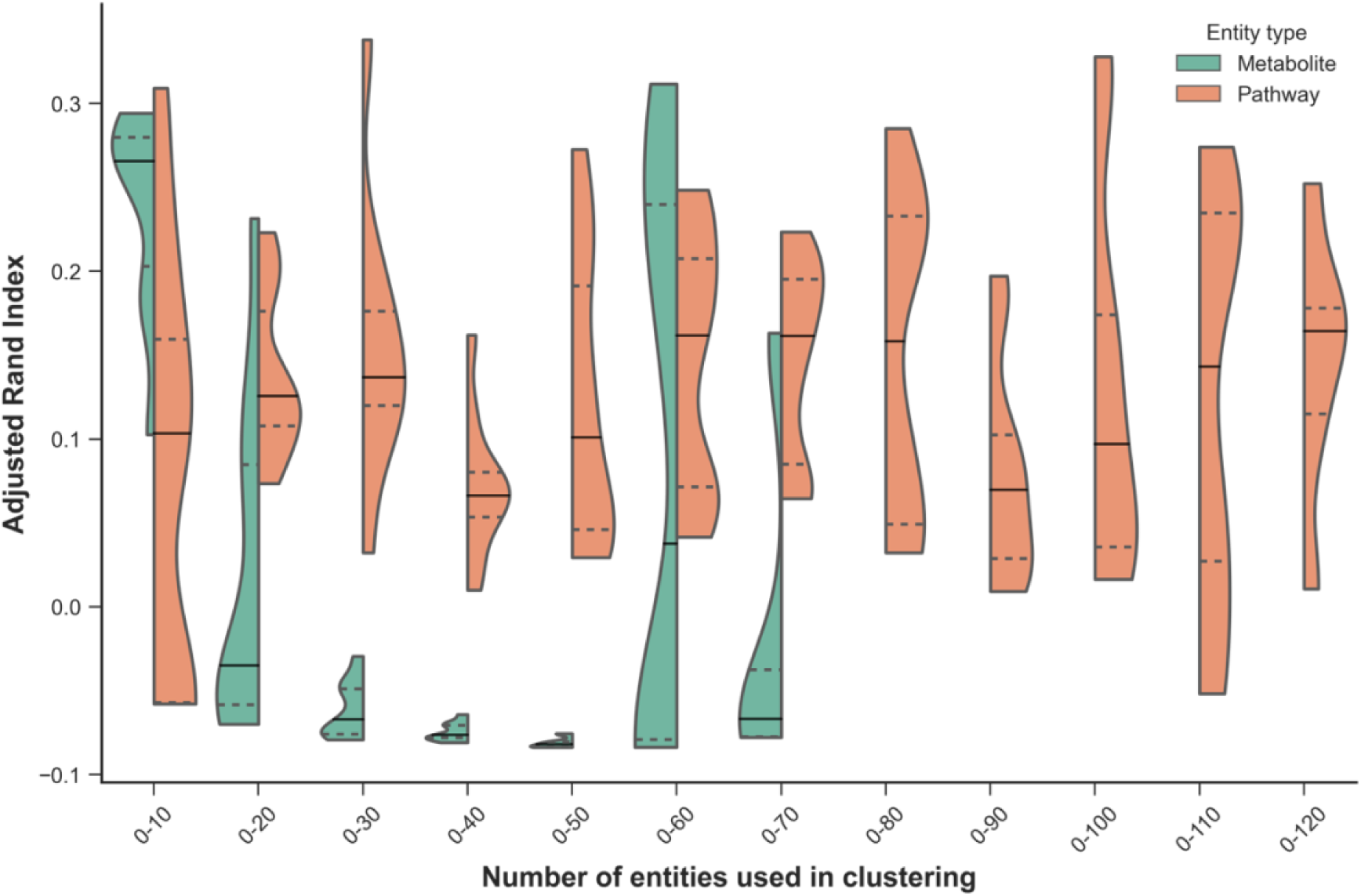
Clustering performance at the metabolite and pathway levels (kPCA method) for different selection thresholds. Clustering performance is quantified using the Adjusted Rand Index (y-axis). The number of entities used to perform the clustering are shown on the x-axis, and only entities significant at q ≤ 0.05 were included in the analysis. A total of 69 metabolites and 116 pathways were significant and are shown in the violin plots. Entities were ranked by q-value and only those within each selection threshold are used for each comparison. Violin plots show the distribution of ARI values achieved using the cumulative thresholds (e.g. the top 10 pathways). Solid lines represent the median ARI, whereas dashed lines represent the upper and lower quartiles. The violin plot range is truncated to the range of the data.

**Fig 7:**
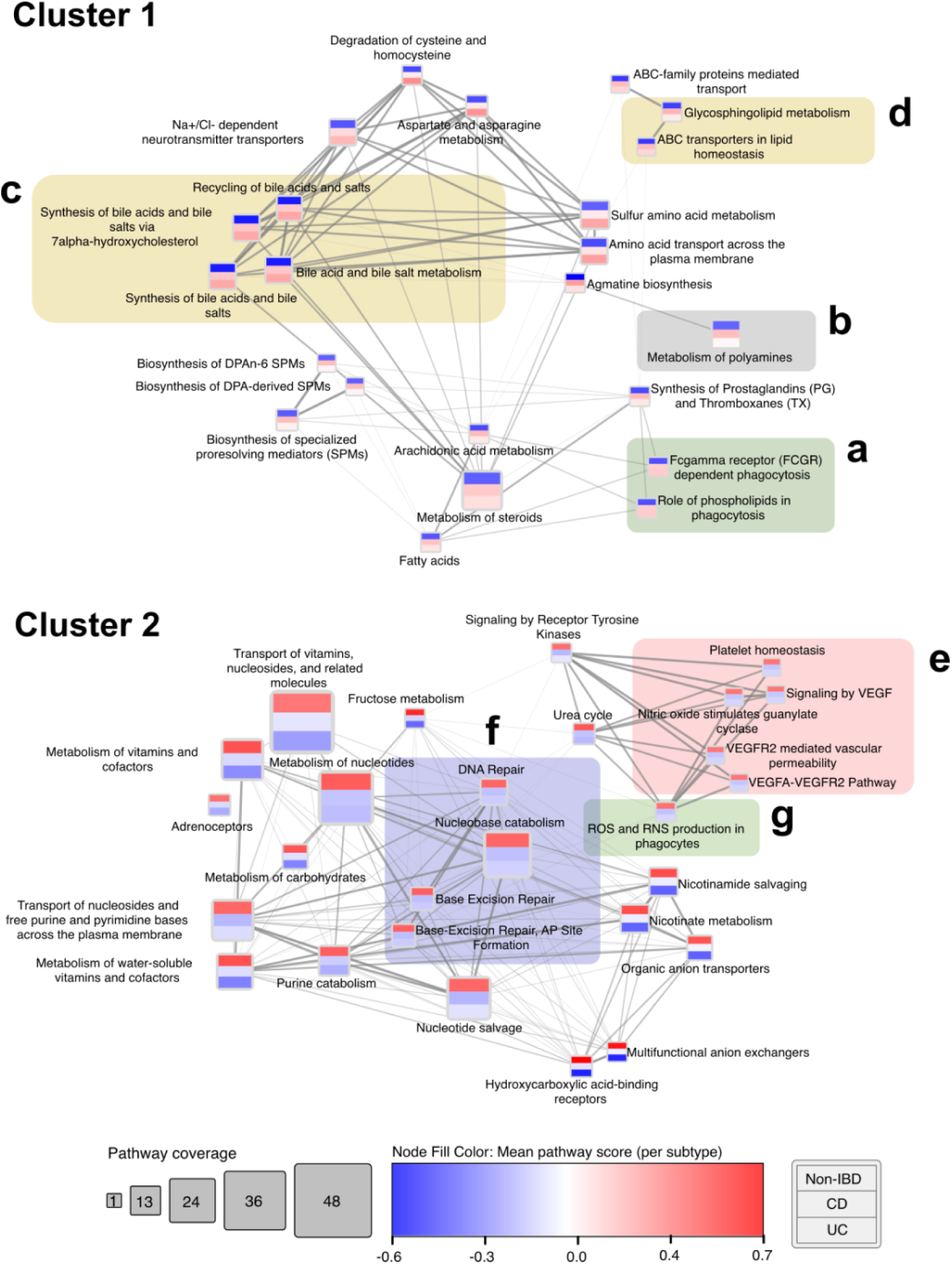
Pathway cluster network derived using hierarchical clustering on IBD pathway scores (kPCA, top 50 pathways). Coloured bands in each node represent mean pathway score within each subtype (from top to bottom: non-IBD, CD, UC). Edge weight represents Spearman correlation between pathway scores. Only edges with Spearman’s rank correlation coefficient (ρ) ≥ 0.4 are displayed. Shaded sub-clusters depict pathways consistent with current IBD literature and are discussed in the text. Shading of subclusters corresponds to Reactome pathway parent category.

The networks shown in Fig. 7 represent two distinct groups of pathways that discriminate between IBD and non-IBD samples. An edge-weighted spring-embedded layout was applied to the network to visually group nodes by pathway score Spearman correlation. The network nodes are coloured by the average pathway score across the samples in each subtype (from top to bottom: non-IBD, CD, UC). Using the kPCA approach (or clustering/DR-based approaches in general), it is not possible to make a direct link between the sign of the pathway scores obtained and the direction of the pathway enrichment. However, GSEA-based ssPA approaches can be used to determine this, if desired. GSVA was used to determine that cluster 1 of Fig. 7 represents pathways whose metabolites are upregulated in IBD relative to the non-IBD group, whereas cluster 2 represents pathways with metabolites depleted in IBD relative to the non-IBD group.

IBD encompasses a group of diseases broadly characterised by chronic inflammation of the gut, and is widely attributed to a complex interplay of immune dysregulation, dysbiosis of the gut microbiome, and genetic and environmental factors. A sub-cluster of immune-related pathways can be seen in cluster 1 (Fig. 7a), highlighting the role of phospholipids and Fc-gamma receptors in phagocytosis, which is concordant with findings in current literature [26,27]. Changes in phospholipid metabolism are strongly implicated in IBD pathology, particularly in degradation of the intestinal epithelial mucus layer [27]. Another pathway associated with maintaining the integrity of the gut mucosal barrier highlighted in this cluster is ‘Metabolism of polyamines’ (Fig. 7b), which corroborates the work of Weiss et al., 2004 [28], who found increased levels of the polyamine spermidine alongside decreased levels of spermine in IBD patients. One major benefit of ssPA is that it enables comparison of pathway scores across multiple study groups. Using the ‘Metabolism of polyamines’ pathway as an example, we note that enrichment of the pathway is greater in CD patients than it is in UC patients, relative to non-IBD patients. A large sub-cluster of differentially active pathways relates to bile acid and salt metabolism (Fig. 7c), consistent with alterations in primary (e.g. cholate) and secondary bile acid metabolism in IBD observed by Lloyd-Price et al., 2019, likely attributable to changes in gut microbial composition, as microbes directly modulate bile acids [29]. Other notable pathways enriched in cluster 1 (particularly in CD patients) include ‘Glycosphingolipid metabolism’ and ‘ABC transporters in lipid homeostasis’ (Fig. 7d), which are congruous with the known dysregulation of lipid metabolism in IBD coupled with inflammation [30–32].

In the second cluster, which consists of pathways whose metabolites are depleted in IBD, a prominent sub-cluster containing pathways related to platelet homeostasis, VEGF, and nitric oxide (NO) signalling is visible (Fig. 7e). It is widely recognised that platelet abnormalities are associated with IBD, alongside complications such as microvascular thrombosis and thromboembolism [33]. The reduction in NO signalling is consistent with findings of reduced NO-mediated vasodilation found in IBD patients [34], which could further contribute to thrombosis. This cluster of pathways focused around VEGF signalling is one example that was not detected as significantly enriched using ORA, but was detected amongst the top pathways using ssPA (see Table S3 for ORA results).

Another large sub-cluster within cluster 2 focuses on DNA repair processes, including base-excision repair (Fig. 7f). This subcluster can be linked to another significant pathway, ‘ROS and RNS production in phagocytes’ (Fig. 7g), as reactive oxygen species (ROS) contribute to DNA damage, which in turn induce DNA repair mechanisms [35,36]. The negative enrichment of these pathways may allude to potential defects or reduction in DNA repair processes in IBD.

We also trained a random forest (RF) classifier on the IBD dataset, which based on 5-fold repeated stratified cross-validation achieved comparable AUC using kPCA scores (*μ* = 0.861, *σ* = 0.042) as did the classifier trained on the pathway-annotated metabolites (*μ* = 0.865, *σ* = 0.040). Pathway importances were calculated by permuting each of the features individually, and ranked by the mean decrease in AUC. Many of the top 50 pathways ranked by the RF classifier are shared with those in the networks in Fig. 7 (see Table S4).

Using the same approach, a pathway-based correlation network was created using the Su et al., 2020dataset (Fig. S6). The use of ssPA shows a clear correlation between the WHO status of the samples, corresponding to COVID severity, and the enrichment level of the top 30 pathways shown in the network. Both case studies highlight the value of ssPA methods in quantifying pathway enrichment at an individual sample level, enabling direct comparison of multiple sample sub-groups, as well as in the case of the IBD network, identifying a cluster of enriched pathways related to VEGF/NO signalling that were not identified by the conventional method ORA. Taken together, these case studies demonstrate a small subset of the downstream analyses possible using pathway scores, but highlight the benefits in interpretability and predictive robustness they can achieve.

## Discussion

Single-sample PA methods have continued to gain traction in recent literature as a way of examine pathway signatures at an individual sample level [19,20,37,38]. Unlike conventional PA approaches, which usually compare two groups of samples, ssPA approaches enable researchers to dissect the heterogeneity of complex diseases or response to treatments at an individual level, facilitating advances in precision medicine. Within the transcriptomics field, continuous advances to ssPA methodologies are being proposed, alongside demonstrated applications on gene-expression data [19,37–39]. Despite the importance of metabolic pathways in disease and drug-response, there exists very little literature surveying the applicability of ssPA approaches to metabolomics data [40]. The present study was designed to address this gap by providing a critical evaluation of the most widely-used ssPA methods and their performance, robustness, and applicability to metabolomics data.

Using semi-synthetic metabolomics data, our benchmarking procedure evaluated several properties of ssPA methods: recall, precision, AUC, and the ability to rank enriched pathways highly. By varying simulation parameters such as effect size and signal strength, we were able to ascertain how the methods performed under these more realistic scenarios. Applied to transcriptomics data, GSVA is often a top performer, particularly in gene-set recall [15,19]. This finding is consistent with our results, where GSVA was found to have higher recall than all other ssPA methods across lower effect sizes, and at higher effect sizes had similar recall to ssGSEA. GSEA-based methods may provide more power at lower effect sizes as they calculate scores as a function of the compounds inside and outside of the pathway, testing a competitive null hypothesis, whereas all other methods benchmarked (besides z-score) calculate scores based only on the compounds within a pathway itself. When effect sizes become larger however, the recall of GSVA was found to be very similar to that of ssGSEA and z-score. In contrast, when identifying correctly the enriched pathways (precision), GSVA was one of the lowest performing methods, as opposed to clustering and DR based methods ssClustPA, kPCA and SVD. In general therefore, it can be inferred that GSEA/z-score-based methods offer higher recall (most evident at lower effect sizes), whereas clustering/DR-based methods offer improvements in precision, particularly at higher effect sizes, for metabolomics data. In general we suggest the use of clustering/DR-based methods for datasets with moderate-high effect sizes, and GSVA for datasets with lower effect sizes.

We offer two novel methods for ssPA: ssClustPA and kPCA, which can theoretically be applied to any omics datatype. Leveraging fundamental machine learning concepts of clustering and kernels respectively, both ssClustPA and kPCA have been designed to make use of these unsupervised approaches to discriminate between samples at the pathway level. ssClustPA is particularly advantageous when there is clear structure to the underlying data, exploiting this to provide more precise pathway scores. The RBF kernel used in the kPCA method allows complex non-linear structure within the data to be modelled and projected into a subspace where datapoints become more linearly separable, and hence more likely to provide precise pathway scores, which would otherwise not be feasible using linear separation methods such as PCA or SVD. Applied to metabolomics data, these methods have been shown to offer advances in precision compared to established ssPA methods. Using metabolomics data from IBD samples, Click or tap here to enter text.we employed kPCA to generate pathway scores, and created an IBD subtype-specific pathway network using this data.

Although the objective of this work was not to ascertain whether metabolite or pathway level data yields better performance in downstream analyses, a key question we were able to partially address was whether pathway-level data exhibits greater predictive robustness than its metabolite-level counterpart [18]. Using the IBD dataset, improved clustering performance was observed when using kPCA pathway scores as opposed to individual metabolites. Not only was the maximum ARI achievable higher using the pathway scores, but the range of ARI scores obtained using a variety of different thresholds used to select which entities to include in the clustering was consistently higher using pathway scores than metabolites. This observation supports the hypothesis that in some cases pathway scores can provide improved robustness to noise, one example of which is the number of pathways used as features in downstream analyses.

We have undertaken the first comprehensive benchmark into ssPA methods for metabolomics data. The insights gained from this study may provide guidance to practitioners in the field for selecting an appropriate ssPA method, as well as highlighting the broad range of applications for downstream analyses using pathway scores. The semi-synthetic simulation approach we have outlined is not only applicable to metabolomics data, but can be used to evaluate a multitude of PA methods across various omics datatypes. Although we used a single pathway resource, Reactome, in both the benchmarking and application sections of this work, our approach is easily extensible to other databases e.g. KEGG [41] or MetaCyc [42] and we do not expect our results to be highly dependent on the pathway database used. A class of methods we have not included in this work are topology-based ssPA methods [7]. Further work is required to determine how such methods compare to non-topology based alternatives, and whether the inclusion of topological information improves performance notwithstanding the low coverage of pathways observed in metabolomics data. Overall, ssPA methods show a great deal of promise both for analysing and interpreting metabolomics data, in addition to the prospect of integrating metabolomics with other omics datatypes.

## Acknowledgements

The authors gratefully acknowledge Fabien Jourdan, Nathalie Poupin, Clément Frainay, and Juliette Cooke for insightful and stimulating discussions regarding this work and for critical reading of the manuscript.

## Data availability

The COVID19 dataset from [22] is available to download from https://data.mendeley.com/datasets/tzydswhhb5/5 (Table S1). The IBD dataset from [23] is available to download from https://www.hmpdacc.org/ihmp or at MetabolomicsWorkbench (PR000639).

## Code availability

Python scripts used to produce the results of this work are publicly available at https://github.com/cwieder/sspa-in-metabolomics. The sspa Python package is available to download from https://pypi.org/project/sspa/ and the source code as well as documentation is available at https://github.com/cwieder/py-ssPA.

## Supporting information

**Table S1:**
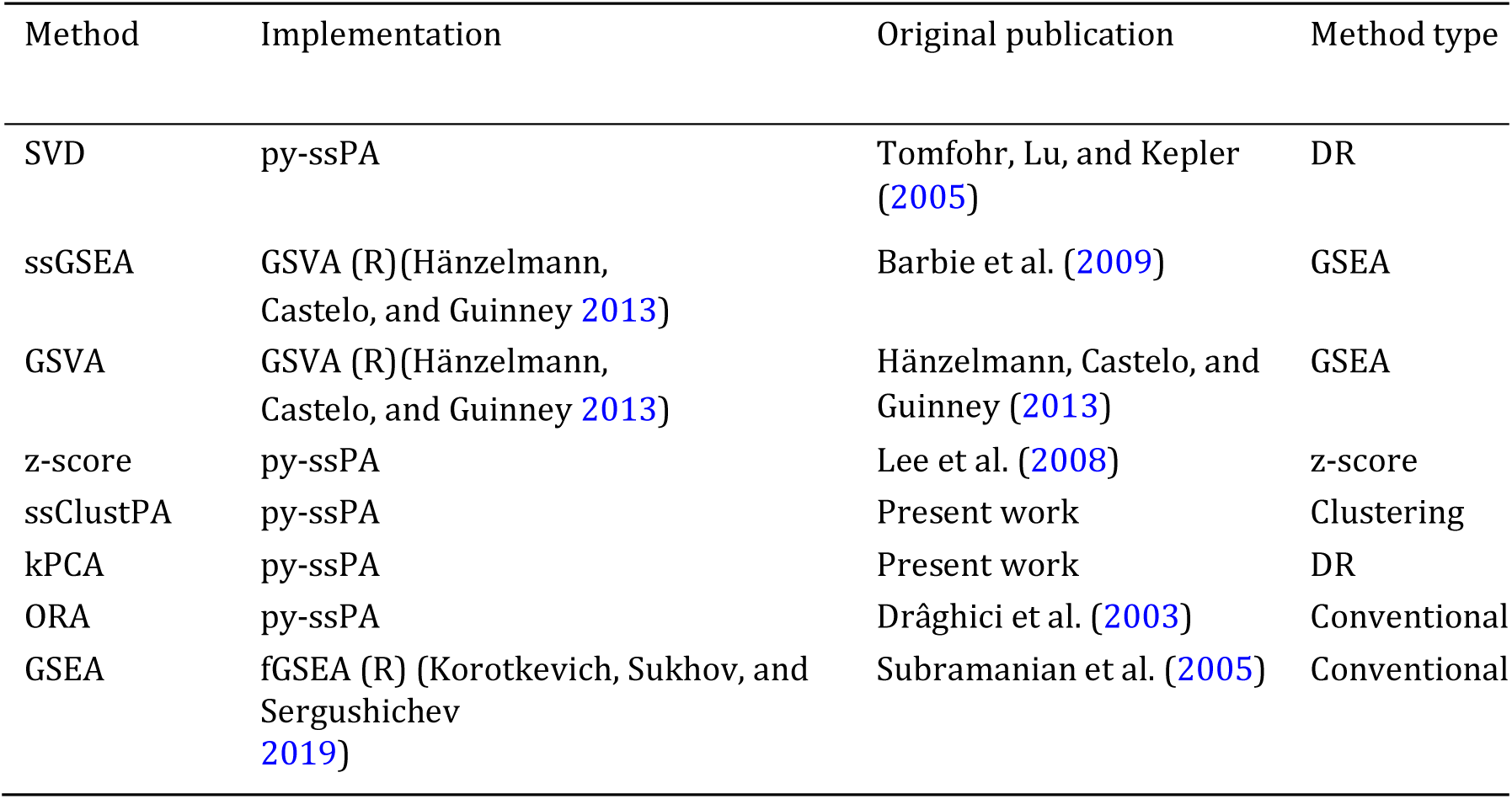
Method implementation details. Methods are classified into categories based on the underlying algorithm used for scoring. Conventional pathway analysis methods i.e. non-single sample are denoted as “Conventional”.

**Fig S1:**
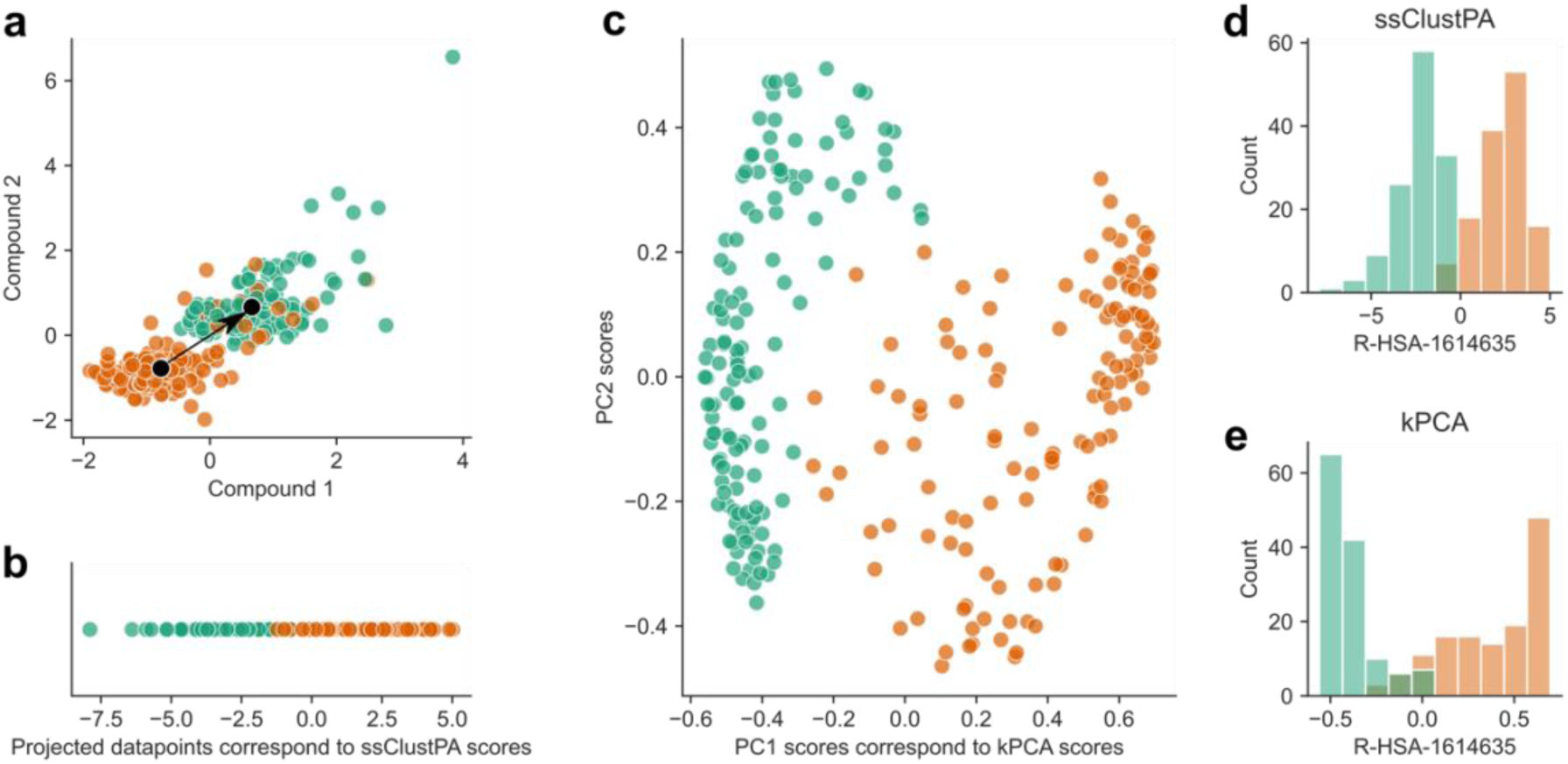
Overview of novel methods ssClustPA and kPCA for a single pathway example using simulated data based on the COVID-19 study from. [22]. Different coloured points are used to distinguish between case and control groups. a) ssClustPA overview. Scatterplot shows datapoints from two dimensions (compounds) of the pathway matrix Z_R–HSA–1614635_ containing all samples but only those metabolites present in R-HSA-1614635. Black dots represent cluster centroids identified by k-means. Scores are calculated by computing the unit vector between the two centroids (black arrow) and projecting this onto the original datapoints. b) ssClustPA scores. Datapoints correspond to those in a), shown after projection onto the unit vector between the two cluster centroids. c) kPCA overview. Scatterplot of kPCA scores (x-axis = PC1 and y-axis = PC2). PC1 scores correspond directly to pathway scores. d, e) Distribution of pathway scores across samples calculated using ssClustPA and kPCA respectively for pathway R-HSA-1614635.

**Fig S2:**
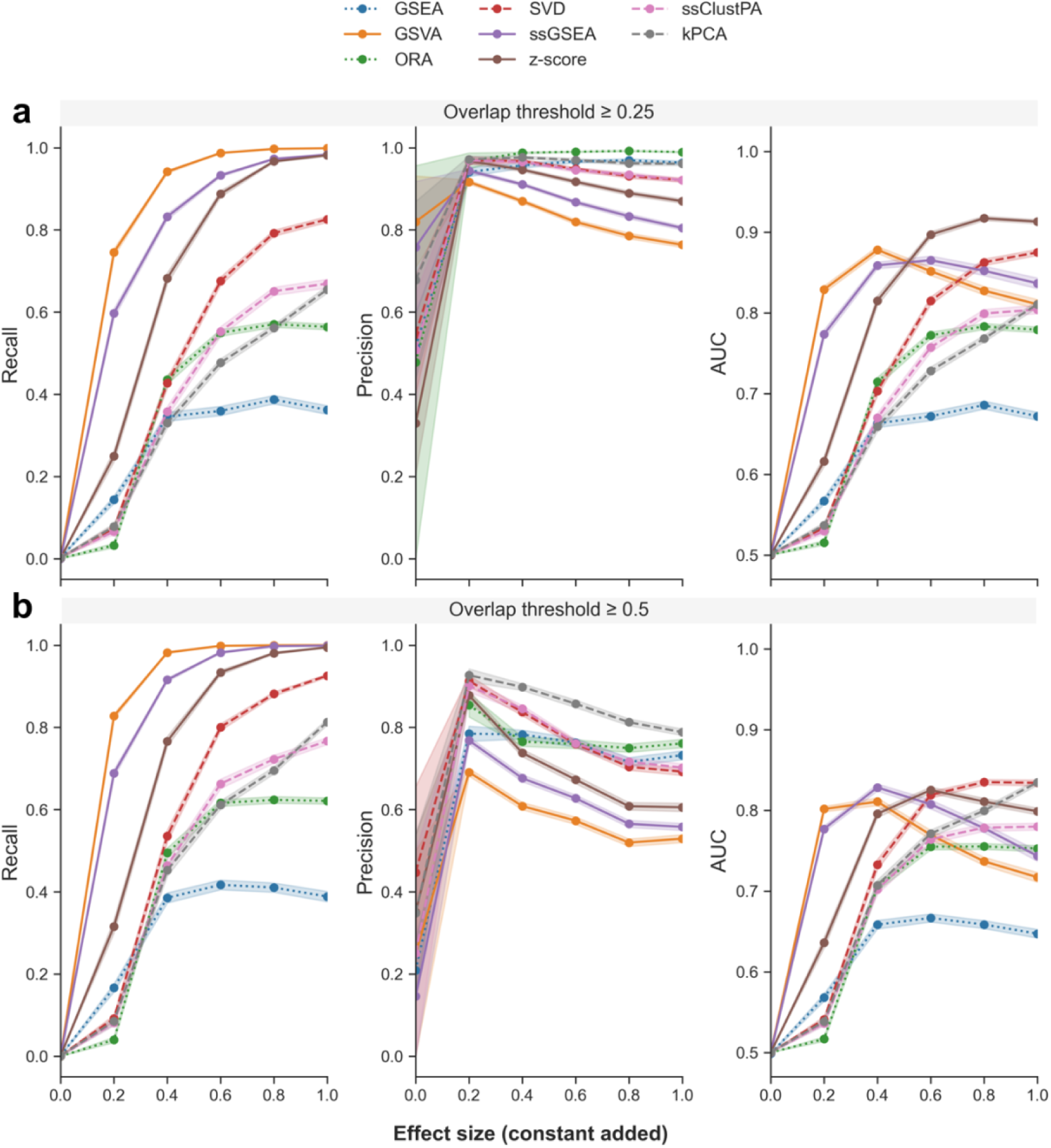
Performance of ssPA methods using the full pathway set (including redundancy) based on 3 randomly enriched pathways, derived using semi-synthetic data based on the IBD dataset. (a) top panel OC ≥ 0.25, (b) bottom panel OC ≥ 0.5. All compounds in enriched pathways have identical effect size. Points show performance metrics averaged across 200 iterations. Shaded intervals represent average SEM. Dotted lines represent conventional pathway analysis methods, and dashed lines represent clustering/dimensionality-reduction based methods.

**Fig S3:**
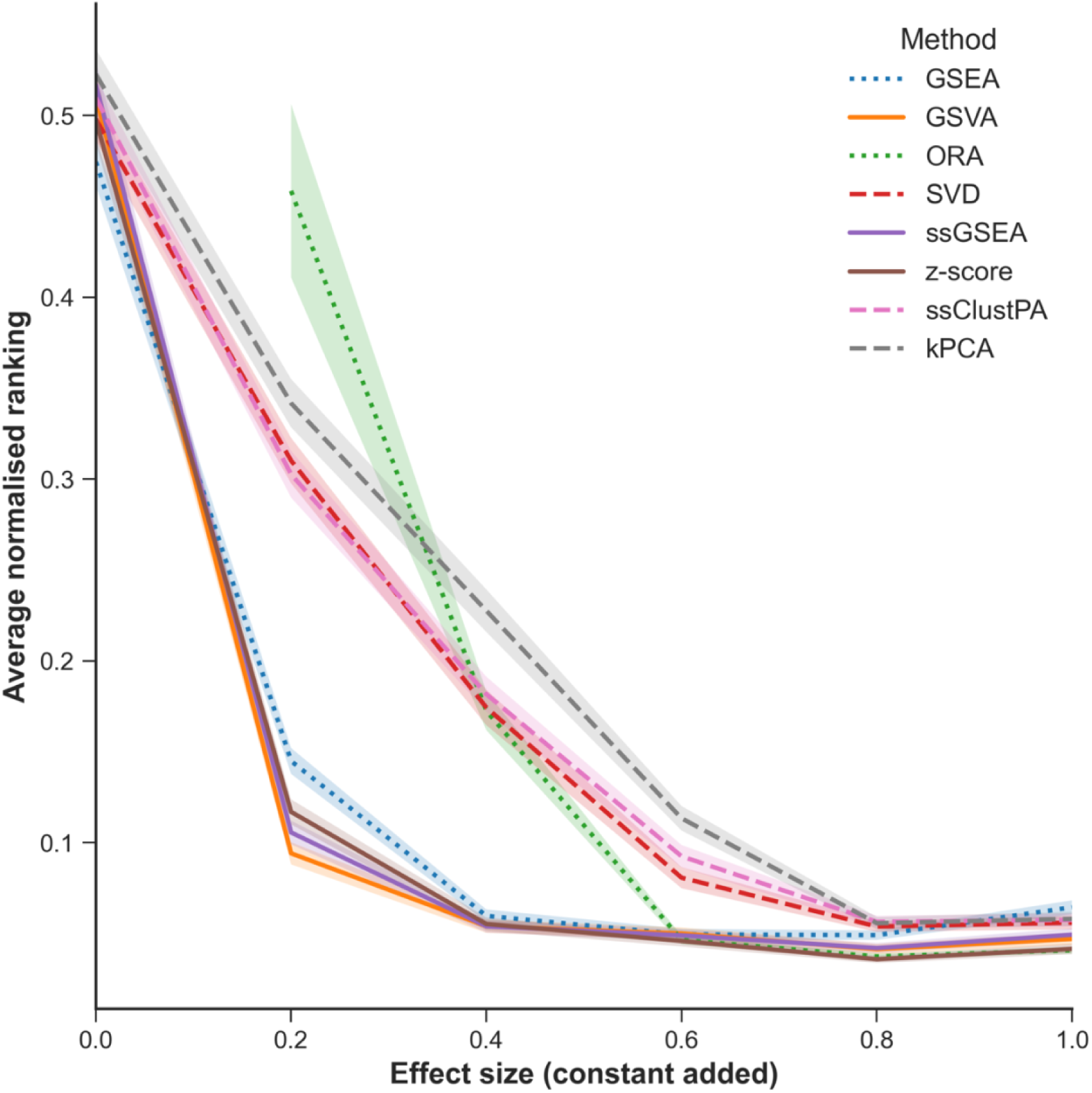
ssPA method ability to rank highly the 3 randomly enriched pathways using the full set of Reactome pathways. Average normalised ranking of 3 randomly selected enriched pathways shown on the y-axis. Shaded intervals represent standard error on the mean averaged over 200 iterations. Dotted lines represent conventional pathway analysis methods, and dashed lines represent clustering/dimensionality-reduction based methods.

**Fig S4:**
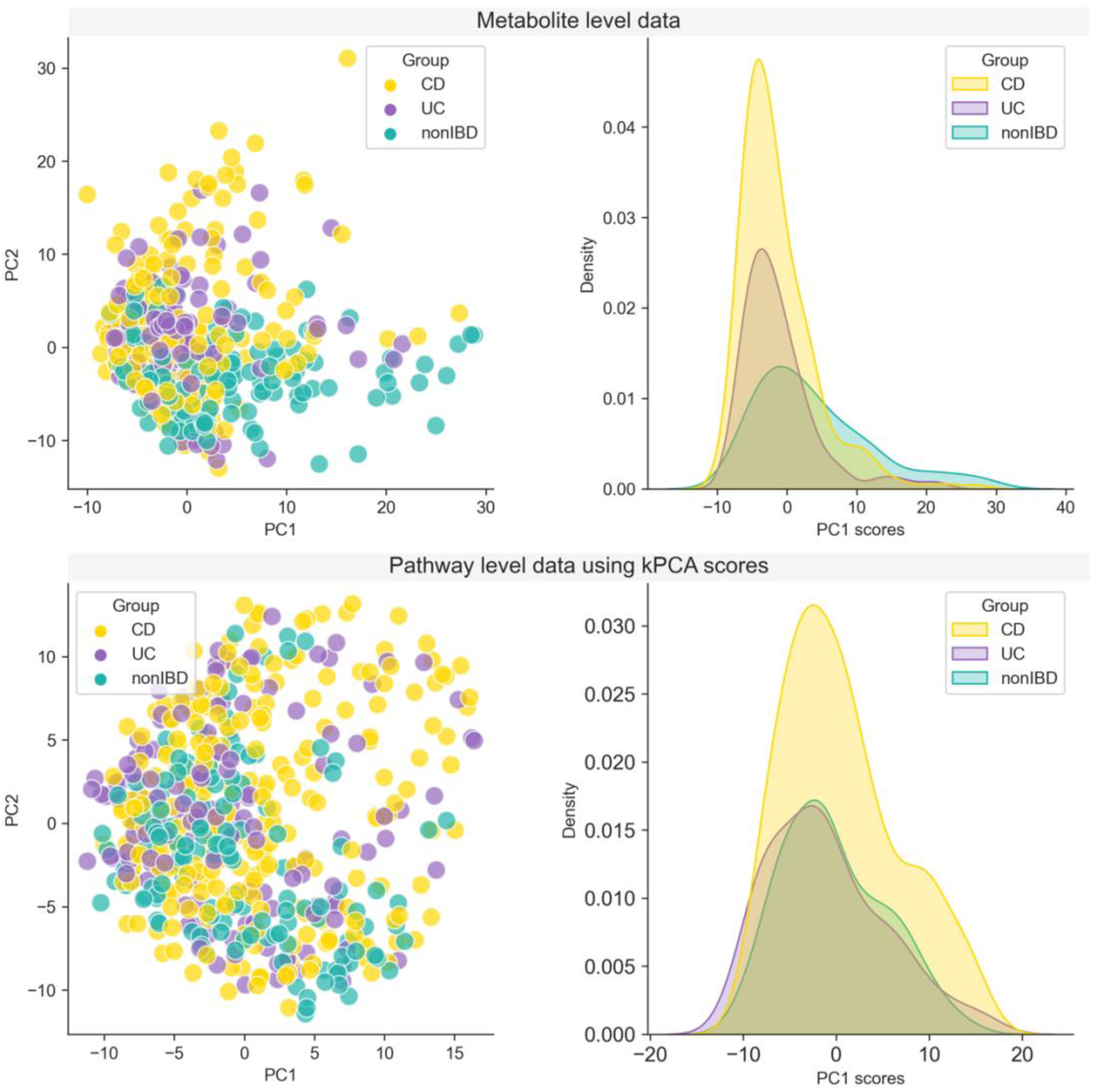
PCA scatter plots and density plots of PC1 scores obtained using the IBD data at the metabolite (upper panels) and pathway level using kPCA (lower panels).

**Fig S5:**
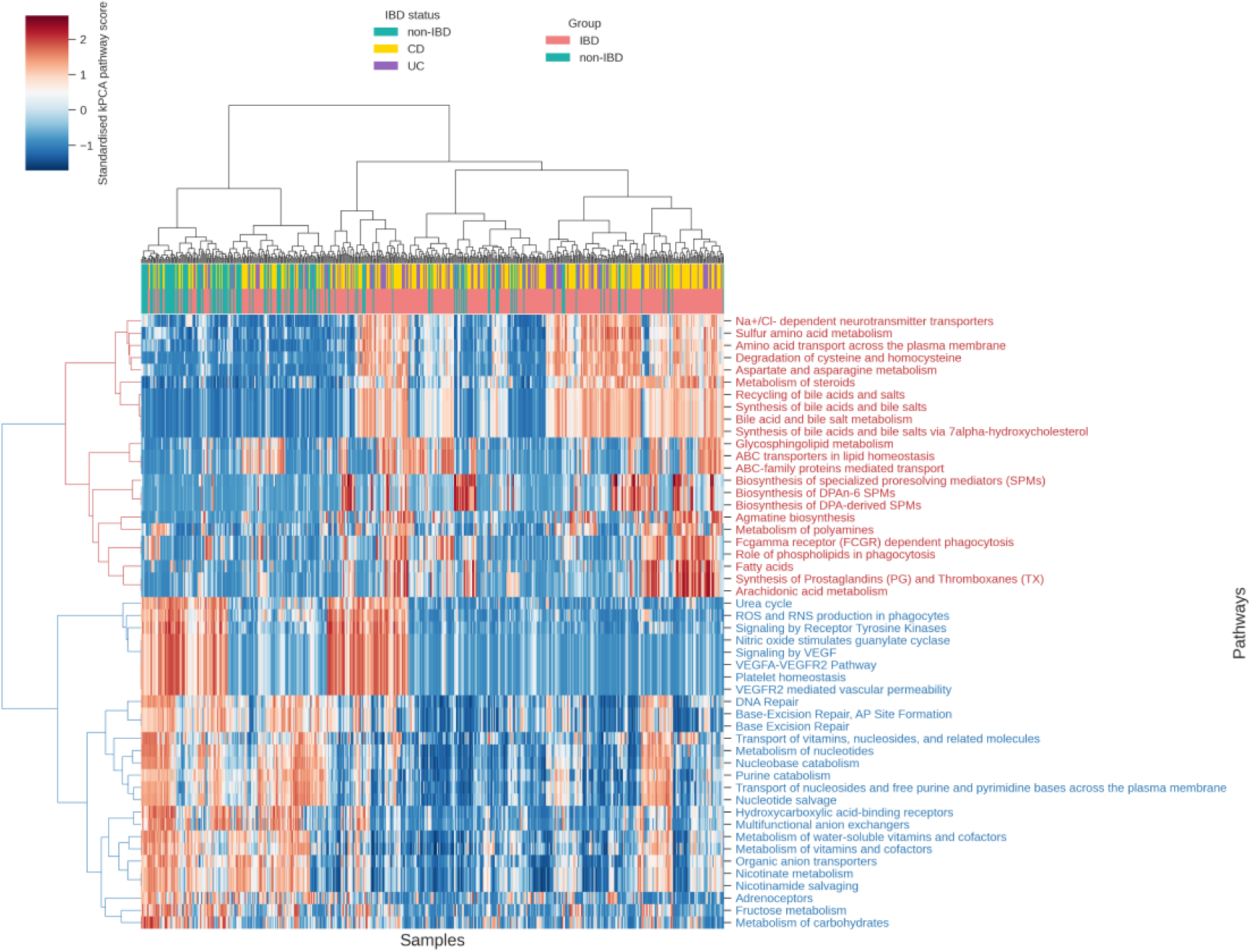
Clustered heatmap of IBD data transformed to pathway scores using the kPCA method. Top 50 pathways are used for clustering (performed on Euclidean distances using Ward linkage).

**Fig S6:**
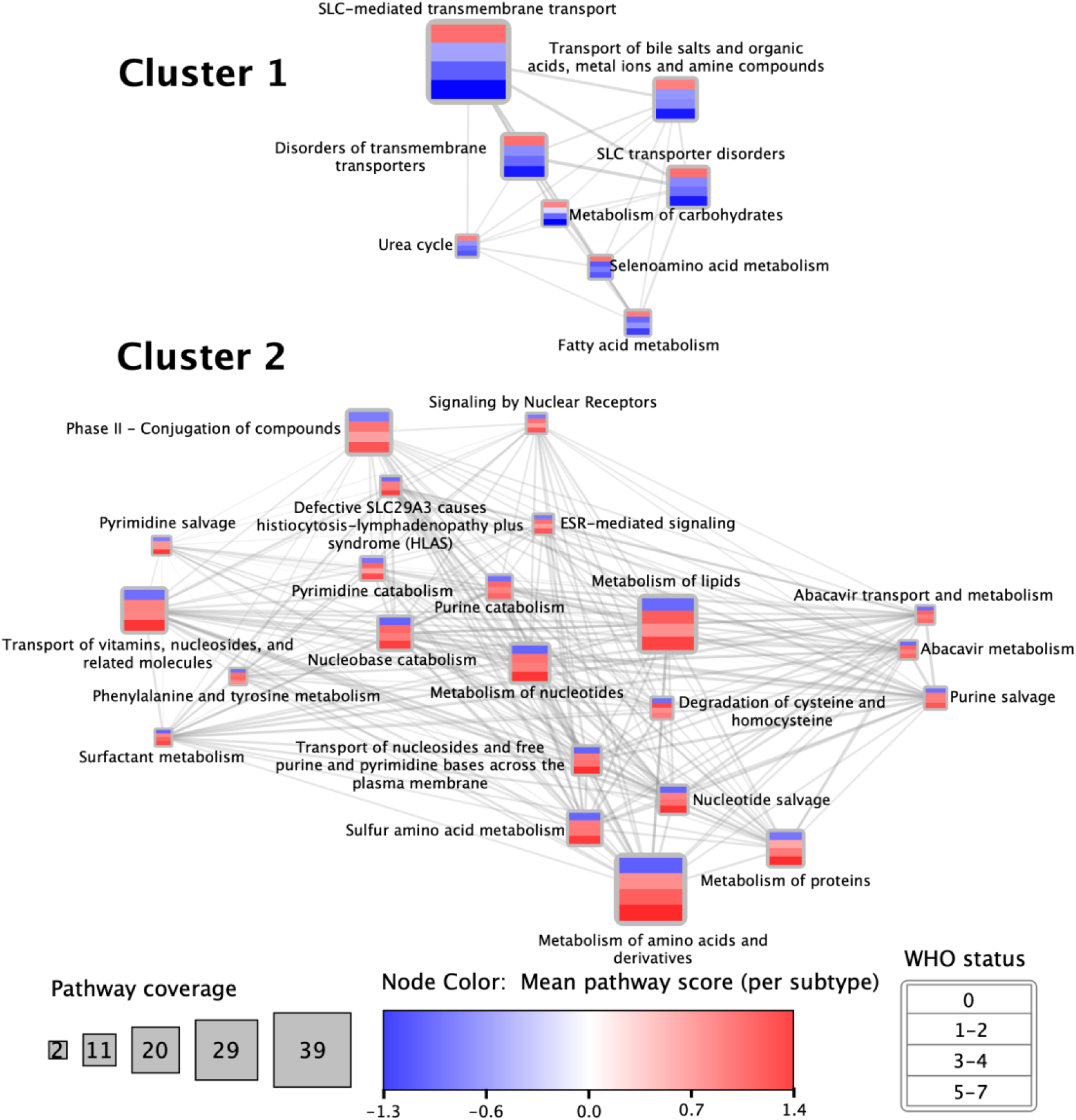
Pathway clusters derived using hierarchical clustering on COVID dataset transformed to pathway scores using kPCA (top 30 pathways). Node fill colour represents mean pathway score across each WHO status level. Edge weight represents Spearman correlation between pathway scores. Only edges with *ρ* ≥ 0.4 are shown.

**Table S2:** Runtimes of ssPA methods, alongside GSEA* for comparison to conventional PA methods (average across 10 iterations). *fGSEA implementation used [43].

| Method | Mean wall time (seconds) |
| --- | --- |
| SVD | 0.43 |
| ssGSEA | 9.62 |
| GSVA | 3.32 |
| z-score | 0.28 |
| ssClustPA | 9.54 |
| kPCA | 2.10 |
| GSEA | 5.10 |

**Table S3:** Over-representation analysis results from IBD dataset.

| Pathway ID | Hits | P-value | P-adjust | Pathway name |
| --- | --- | --- | --- | --- |
| R-HSA-73884 | 4/4 | 0.06 | 0.985 | Base Excision Repair |
| R-HSA-73929 | 4/4 | 0.06 | 0.985 | Base-Excision Repair, AP Site Formation |
| R-HSA-83936 | 10/14 | 0.082 | 0.985 | Transport of nucleosides and free purine and pyrimidine bases across the plasma membrane |
| R-HSA-15869 | 14/21 | 0.084 | 0.985 | Metabolism of nucleotides |
| R-HSA-6798163 | 3/3 | 0.122 | 0.985 | Choline catabolism |
| R-HSA-74259 | 6/8 | 0.139 | 0.985 | Purine catabolism |
| R-HSA-8956321 | 10/15 | 0.141 | 0.985 | Nucleotide salvage |
| R-HSA-425397 | 16/26 | 0.148 | 0.985 | Transport of vitamins, nucleosides, and related molecules |
| R-HSA-8956319 | 11/17 | 0.155 | 0.985 | Nucleobase catabolism |
| R-HSA-159418 | 4/5 | 0.183 | 0.985 | Recycling of bile acids and salts |
| R-HSA-112310 | 8/12 | 0.185 | 0.985 | Neurotransmitter release cycle |
| R-HSA-74217 | 5/7 | 0.221 | 0.985 | Purine salvage |
| R-HSA-888590 | 2/2 | 0.247 | 0.985 | GABA synthesis, release, reuptake and degradation |
| R-HSA-2046105 | 2/2 | 0.247 | 0.985 | Linoleic acid (LA) metabolism |
| R-HSA-71240 | 2/2 | 0.247 | 0.985 | Tryptophan catabolism |
| R-HSA-2029485 | 2/2 | 0.247 | 0.985 | Role of phospholipids in phagocytosis |
| R-HSA-422085 | 2/2 | 0.247 | 0.985 | Synthesis, secretion, and deacylation of Ghrelin |
| R-HSA-888593 | 2/2 | 0.247 | 0.985 | Reuptake of GABA |
| R-HSA-2142753 | 2/2 | 0.247 | 0.985 | Arachidonic acid metabolism |
| R-HSA-2162123 | 2/2 | 0.247 | 0.985 | Synthesis of Prostaglandins (PG) and Thromboxanes (TX) |
| R-HSA-193807 | 2/2 | 0.247 | 0.985 | Synthesis of bile acids and bile salts via 27-hydroxycholesterol |
| R-HSA-1222556 | 2/2 | 0.247 | 0.985 | ROS and RNS production in phagocytes |
| R-HSA-2029480 | 2/2 | 0.247 | 0.985 | Fcγ receptor (FCGR) dependent phagocytosis |
| R-HSA-444209 | 8/13 | 0.282 | 0.985 | Free fatty acid receptors |
| R-HSA-2046104 | 3/4 | 0.309 | 0.985 | α-linolenic (omega3) and linoleic (omega6) acid metabolism |
| R-HSA-109582 | 3/4 | 0.309 | 0.985 | Hemostasis |
| R-HSA-442660 | 4/6 | 0.339 | 0.985 | Na<sup>+</sup>/Cl<sup>-</sup> dependent neurotransmitter transporters |
| R-HSA-73894 | 4/6 | 0.339 | 0.985 | DNA Repair |
| R-HSA-194068 | 4/6 | 0.339 | 0.985 | Bile acid and bile salt metabolism |
| R-HSA-193368 | 4/6 | 0.339 | 0.985 | Synthesis of bile acids and bile salts via 7α-hydroxycholesterol |
| R-HSA-192105 | 4/6 | 0.339 | 0.985 | Synthesis of bile acids and bile salts |
| R-HSA-71291 | 19/35 | 0.351 | 0.985 | Metabolism of amino acids and derivatives |
| R-HSA-168249 | 5/8 | 0.358 | 0.985 | Innate Immune System |
| R-HSA-168256 | 5/8 | 0.358 | 0.985 | Immune System |
| R-HSA-2980736 | 8/14 | 0.388 | 0.985 | Peptide hormone metabolism |
| R-HSA-112316 | 8/14 | 0.388 | 0.985 | Neuronal System |
| R-HSA-8978868 | 8/14 | 0.388 | 0.985 | Fatty acid metabolism |
| R-HSA-112315 | 8/14 | 0.388 | 0.985 | Transmission across Chemical Synapses |
| R-HSA-5619102 | 11/20 | 0.402 | 0.985 | SLC transporter disorders |
| R-HSA-373076 | 15/28 | 0.414 | 0.985 | Class A/1 (Rhodopsin-like receptors) |
| R-HSA-425407 | 25/48 | 0.427 | 0.985 | SLC-mediated transmembrane transport |
| R-HSA-556833 | 16/31 | 0.492 | 0.985 | Metabolism of lipids |
| R-HSA-416476 | 10/19 | 0.494 | 0.985 | G α (q) signalling events |
| R-HSA-1614635 | 4/7 | 0.496 | 0.985 | Sulfur amino acid metabolism |
| R-HSA-418555 | 4/7 | 0.496 | 0.985 | G α (s) signalling events |
| R-HSA-9664433 | 4/7 | 0.496 | 0.985 | Leishmania parasite growth and survival |
| R-HSA-9662851 | 4/7 | 0.496 | 0.985 | Anti-inflammatory response favouring Leishmania parasite infection |
| R-HSA-196807 | 4/7 | 0.496 | 0.985 | Nicotinate metabolism |
| R-HSA-197264 | 4/7 | 0.496 | 0.985 | Nicotinamide salvaging |
| R-HSA-5619108 | 4/7 | 0.496 | 0.985 | Defective SLC27A4 causes ichthyosis prematurity syndrome (IPS) |
| R-HSA-9658195 | 4/7 | 0.496 | 0.985 | Leishmania infection |
| R-HSA-804914 | 4/7 | 0.496 | 0.985 | Transport of fatty acids |
| R-HSA-73614 | 4/7 | 0.496 | 0.985 | Pyrimidine salvage |
| R-HSA-9660821 | 4/7 | 0.496 | 0.985 | ADORA2B mediated anti-inflammatory cytokines production |
| R-HSA-73621 | 5/9 | 0.496 | 0.985 | Pyrimidine catabolism |
| R-HSA-390696 | 2/3 | 0.497 | 0.985 | Adrenoceptors |
| R-HSA-70635 | 2/3 | 0.497 | 0.985 | Urea cycle |
| R-HSA-6806667 | 2/3 | 0.497 | 0.985 | Metabolism of fat-soluble vitamins |
| R-HSA-6814848 | 2/3 | 0.497 | 0.985 | Glycerophospholipid catabolism |
| R-HSA-76002 | 2/3 | 0.497 | 0.985 | Platelet activation, signaling and aggregation |
| R-HSA-8963693 | 2/3 | 0.497 | 0.985 | Aspartate and asparagine metabolism |
| R-HSA-9018678 | 2/3 | 0.497 | 0.985 | Biosynthesis of specialized proresolving mediators (SPMs) |

**Table S4:** Top 50 features in random forest model based on IBD data. Features are ranked by mean AUC decrease which is computed by permuting each of the features individually.

| Pathway ID | Mean AUC decrease | Pathway name |
| --- | --- | --- |
| R-HSA-3296197 | 0.00979 | Hydroxycarboxylic acid-binding receptors |
| R-HSA-196854 | 0.00391 | Metabolism of vitamins and cofactors |
| R-HSA-5652084 | 0.00333 | Fructose metabolism |
| R-HSA-9018683 | 0.00263 | Biosynthesis of DPA-derived SPMs |
| R-HSA-428643 | 0.00262 | Organic anion transporters |
| R-HSA-9658195 | 0.00194 | Leishmania infection |
| R-HSA-211935 | 0.00172 | Fatty acids |
| R-HSA-9018678 | 0.00166 | Biosynthesis of specialized proresolving mediators (SPMs) |
| R-HSA-177128 | 0.00148 | Conjugation of salicylate with glycine |
| R-HSA-1614558 | 0.00147 | Degradation of cysteine and homocysteine |
| R-HSA-156587 | 0.00146 | Amino Acid conjugation |
| R-HSA-156581 | 0.00135 | Methylation |
| R-HSA-192105 | 0.00126 | Synthesis of bile acids and bile salts |
| R-HSA-159424 | 0.00119 | Conjugation of carboxylic acids |
| R-HSA-193368 | 0.00117 | Synthesis of bile acids and bile salts via 7alpha-hydroxycholesterol |
| R-HSA-427601 | 0.00112 | Multifunctional anion exchangers |
| R-HSA-197264 | 0.00104 | Nicotinamide salvaging |
| R-HSA-75105 | 0.00104 | Fatty acyl-CoA biosynthesis |
| R-HSA-418555 | 0.001 | G alpha (s) signalling events |
| R-HSA-2029485 | 0.00094 | Role of phospholipids in phagocytosis |
| R-HSA-194068 | 0.00084 | Bile acid and bile salt metabolism |
| R-HSA-3134975 | 0.00084 | Regulation of innate immune responses to cytosolic DNA |
| R-HSA-83936 | 0.00084 | Transport of nucleosides and free purine and pyrimidine bases across the plasma membrane |
| R-HSA-209776 | 0.00081 | Metabolism of amine-derived hormones |
| R-HSA-2046105 | 0.0008 | Linoleic acid (LA) metabolism |
| R-HSA-1369062 | 0.00078 | ABC transporters in lipid homeostasis |
| R-HSA-9025106 | 0.00077 | Biosynthesis of DPAn-6 SPMs |
| R-HSA-211859 | 0.00075 | Biological oxidations |
| R-HSA-351143 | 0.00075 | Agmatine biosynthesis |
| R-HSA-425397 | 0.00075 | Transport of vitamins, nucleosides, and related molecules |
| R-HSA-9711123 | 0.00074 | Cellular response to chemical stress |
| R-HSA-9660821 | 0.00072 | ADORA2B mediated anti-inflammatory cytokines production |
| R-HSA-196849 | 0.00072 | Metabolism of water-soluble vitamins and cofactors |
| R-HSA-156580 | 0.0007 | Phase II - Conjugation of compounds |
| R-HSA-390696 | 0.0007 | Adrenoceptors |
| R-HSA-211897 | 0.00067 | Cytochrome P450 - arranged by substrate type |
| R-HSA-189483 | 0.00065 | Heme degradation |
| R-HSA-162582 | 0.00065 | Signal Transduction |
| R-HSA-156584 | 0.00064 | Cytosolic sulfonation of small molecules |
| R-HSA-418346 | 0.00063 | Platelet homeostasis |
| R-HSA-9006934 | 0.00062 | Signaling by Receptor Tyrosine Kinases |
| R-HSA-5218920 | 0.00061 | VEGFR2 mediated vascular permeability |
| R-HSA-420499 | 0.0006 | Class C/3 (Metabotropic glutamate/pheromone receptors) |
| R-HSA-8963743 | 0.00058 | Digestion and absorption |
| R-HSA-9707564 | 0.00058 | Cytoprotection by HMOX1 |
| R-HSA-70921 | 0.00057 | Histidine catabolism |
| R-HSA-15869 | 0.00057 | Metabolism of nucleotides |
| R-HSA-74259 | 0.00056 | Purine catabolism |
| R-HSA-5619102 | 0.00055 | SLC transporter disorders |
| R-HSA-211981 | 0.00054 | Xenobiotics |

